# Unraveling the molecular mechanisms underlying spontaneous multipolar mitosis through CIN-seq

**DOI:** 10.64898/2025.12.24.696383

**Authors:** Pin-Rui Su, Ting-Chun Chou, Maria Teresa López-Cascales, Junfeng Hunag, Tsai-Ying Chen, Sjoukje van Wieren, Chan Li, Cecile Beerens, Li You, Jelle Storteboom, Miao-Ping Chien

**Author notes:** These authors contributed equally (Pin-Rui Su, Ting-Chun Chou). These authors contributed equally (Maria Teresa López-Cascales, Junfeng Hunag, Tsai-Ying Chen). Senior author (Miao-Ping Chien).

## Abstract

Multipolar mitosis, a hallmark of chromosomal instability (CIN), drives tumor heterogeneity and therapy resistance, yet remains difficult to study in live cells due to its rare and dynamic nature. To address this, we developed CIN-seq, a targeted single-cell multiomics method that enables large-scale profiling of rare CIN phenotypes and captures their temporally regulated gene expression. Applying CIN-seq, we investigated viable, spontaneous multipolar mitosis, an abnormal division process that cancer cells can survive. Genomic analysis revealed that this mitosis produces polyploid or chromosomally variable progeny, confirming its role in genomic instability. Aneuploidy of Chromosome 16 was linked to increased tripolar mitosis, a finding validated with CRISPR imaging. Transcriptomic analysis showed activation of the Rho GTPase cycle, which was associated with cytokinesis failure, while PTEN attenuation emerged as a key player of viable multipolar mitosis by promoting cell cycle progression and survival via BCL2L1. We also uncovered a novel link between this phenotype and degranulation-like stress responses, which may contribute to cancer cell adaptation to chromosomal instability. Overall, CIN-seq offers a powerful approach for studying rare, live CIN events at single-cell resolution and reveals new mechanisms by which cancer cells adapt to chromosomal instability.

## 1. Introduction

Multipolar mitosis is a predominant phenotype of chromosomal instability (CIN)^[1]^, a process characterized by ongoing chromosomal mis-segregation, which leads to greater cell-to-cell variability in chromosomal changes^[2]^. Unlike normal bipolar mitosis, where chromosomes from the parent cell segregate into two poles, maintaining a diploid chromosome number, multipolar mitosis occurs when chromosomes mis-segregate into multiple poles. Consequently, this enables cancer cells to develop complex genetic profiles, thereby contributing to tumor evolution through the acquisition of metastatic potential and/or resistance to therapies^[3–5]^. Several pathways have been shown to drive different CIN phenotypes through drug perturbation or genetic modifications, which can expand the population of cells conveniently displaying these atypical phenotypes^[3, 6, 7]^. However, these perturbations overlook the mechanisms driving spontaneous multipolar mitosis in cell subpopulations and often lead to cell apoptosis following abnormal mitosis, which is typically not oncogenic in that sense. Furthermore, it has been shown that multipolar mitoses are often preceded by cytokinesis failure^[8, 9]^, however, the occurrence of failed cytokinesis and subsequent fusion of daughter cells does not necessarily guarantee the formation of multipolar mitosis^[8, 10]^. Cells undergoing spontaneous multipolar mitosis, a phenomenon occurring naturally in tumors, have been shown to produce viable daughter cells capable of completing subsequent mitoses^[8]^. This process can lead to the formation of surviving populations, contributing to tumor progression and potentially leading to therapy resistance over time^[3, 4, 11]^. However, the fundamental mechanisms driving this process remain elusive. While single-cell sequencing enables the profiling of heterogeneous cell subpopulations, including those exhibiting CIN phenotypes such as multipolar mitosis, current state-of-the-art single-cell omics sequencing cannot directly link CIN phenotypes to the profiled genotypes or capture CIN associated temporal RNA expression. The aforementioned challenges have limited the development of cancer therapeutics targeting CIN^[3]^. To this end, we developed CIN-seq, an imaging-guided single-cell sequencing method that enables the selective sequencing of CIN cells in action—specifically, sequencing single cells exhibiting live abnormal mitotic or CIN phenotypes, such as multipolar mitosis. Unlike conventional approaches limited to profiling only a few cells exhibiting specific microscopically observable phenotypes^[7, 12]^, CIN-seq allows for simultaneous profiling of thousands, or even much more, of these rare cells that share the same visually observable phenotype, in a high-throughput manner, as we can screen from a population of 10^5-6^ cells. This advancement allows for a thorough comprehension of the underlying mechanisms behind these aberrant cellular phenomena. Furthermore, CIN-seq enables phenotype-to-genotype linking, meaning that the profiled molecular data are associated with observable traits or behaviors—in this context, aberrant mitosis—and captures CIN-associated temporal RNA expression or newly generated RNA transcripts, thereby fostering the discovery of new insights into CIN.

In this study, using MCF10A mammary cells as a model, CIN-seq provided direct insights into the key mechanisms underlying naturally occurring multipolar mitosis—without the need for drug induction or genetic perturbation. Here we show that aneuploidy in Chromosome 16 is correlated with an increased tripolar mitosis rate. Furthermore, upregulation of the RhoGTPase cycle pathway was found to be correlated with cytokinesis failure, leading to polyploid cell formation; however, this pathway alone was insufficient to induce multipolar mitosis. Whereas PTEN attenuation, coupled with accelerated cell cycle progression, suppression of cell cycle checkpoint pathways, and enhanced BCL2L1 expression, is strongly linked to viable and successful multipolar mitosis.

## 2. Results

### 2.1 Development of CIN-seq for screening, detection and single-cell omics profiling of cell subpopulations undergoing live, spontaneous multipolar mitosis

To effectively screen, identify and isolate rare subpopulations of cells exhibiting live, spontaneous multipolar mitosis, a system enabling large-scale microscopic scanning (∼10^5-6^ cells), real-time image analysis of big imaging data, and immediate single-cell isolation followed by downstream single-cell sequencing is required. CIN-seq (**Figure 1a**) is then introduced to screen 10^5-6^ cells stained with nuclear dyes, using a custom-built ultrawide field-of-view optical (UFO) microscope^[13]^ (**Experimental Section**). CIN-seq implements real-time cell segmentation, ground-truth assisted trainable Weka segmentation (GTWeka)^[14]^ (**Experimental Section, Figure S1**), and instant detection of single cells displaying CIN or abnormal mitotic phenotypes, including tripolar mitosis (the most dominant multipolar mitosis; also in this study), lagging chromosomes and chromatin bridges (**Figure S2**, **Experimental Section**). To detect cells exhibiting tripolar mitosis (referred to as TP cells), we developed a fixed-radius near-neighbor search approach (**Figure 1b; Experimental Section**), where tripolar mitoses are detected based on specific distance thresholds, established from a substantial quantity of cells (**Experimental Section**). Specifically, the detection is based on the distance thresholds between the TP mother cell and TP daughter cells (D_MD_), as well as the distances between all pairs of TP daughter cells (D_DD_) (**Figure 1b, Experimental Section**).

**Figure 1.**
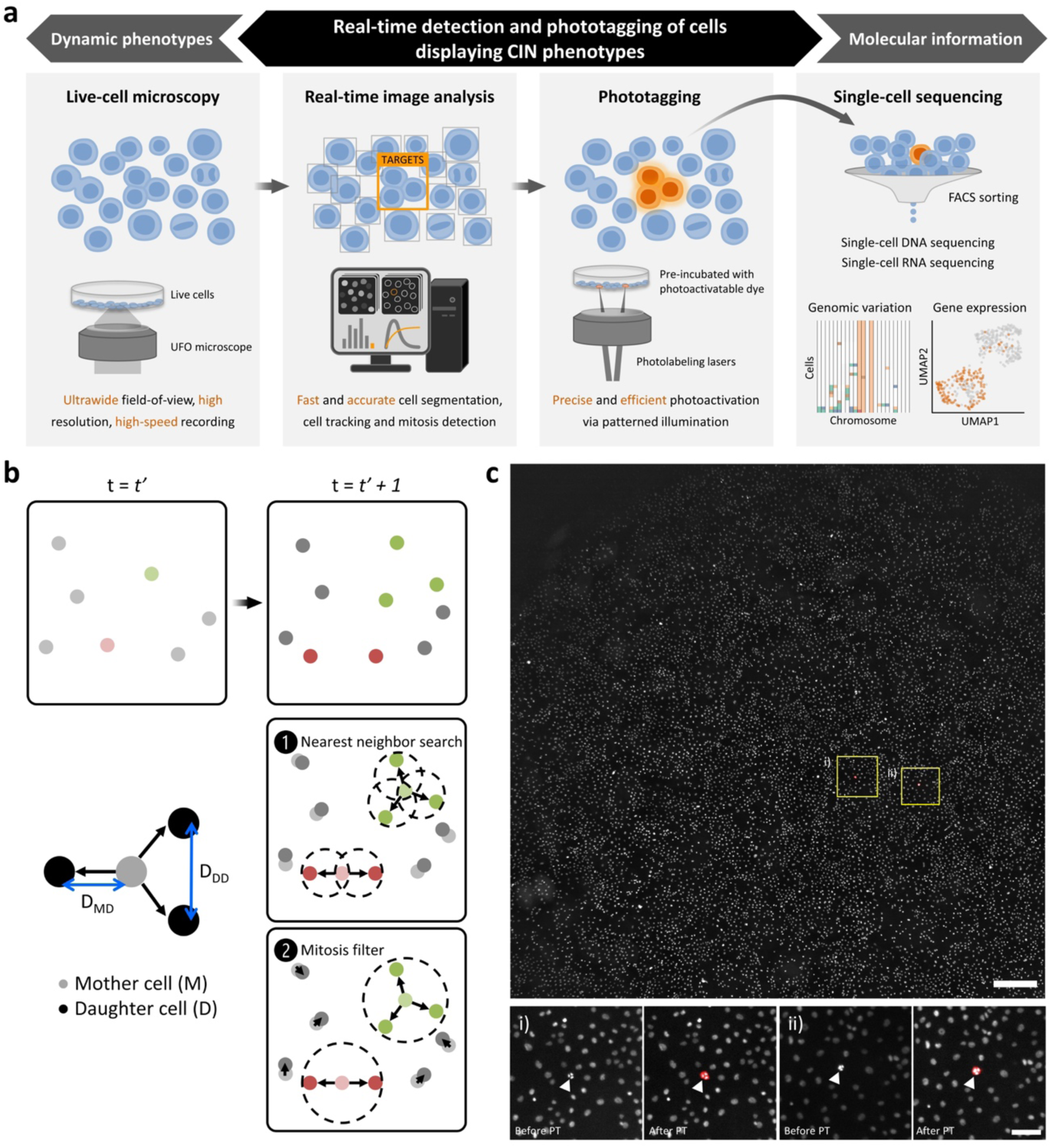
CIN-seq workflow. a) A large quantity of live cells (∼10^5-6^) were imaged through the ultrawide field-of-view optical (UFO) microscope. Subsequently, these images acquired in real-time were instantly processed and analyzed to identify single cells of interest, exhibiting CIN phenotypes within this context. These identified target cells, preincubated with photoactivatable dyes, were immediately and automatically phototagged in real-time. Following this, the cells were detached and subjected to fluorescence activated cell sorting (FACS) to separate phototagged target cells into 384-well plates for downstream single-cell sequencing. b) Mitosis detection is performed using the fixed-radius near neighbor search approach, which relies on specific distance thresholds (mitosis filter) between the TP mother cell and TP daughter cells (D_MD_), and distances between all pairs of TP daughter cells (D_DD_). c) Top panel: A representative image of MCF10A cells, stained with SPY650-DNA nuclear dyes, captured in a field-of-view (FOV) of the UFO microscope. Within this FOV, two instances of tripolar mitoses were detected (i and ii). The scale bar denotes 500 µm. Bottom panel: Zoomed-in imaged showed these two sets of TP cells (white arrows) both before and after phototagging (PT, red). The scale bar denotes 100 µm.

This fixed-radius near-neighbor search method has proven highly effective, accurately detecting TP mitosis with a F1 score of 0.91, which was achieved within a fast processing time of 10 seconds per frame, encompassing 25,000 cells. This high-throughput detection of mitotic events or CIN phenotypes across a substantial cell population enables effective observation of rare events and later facilitates the isolation of rare cells exhibiting CIN phenotypes. Subsequently, within the CIN-seq pipeline, the detected TP cells were automatically phototagged (**Figure 1c**), during real-time time-lapse recordings, acquired within 2-5 hours (**Experimental Section**). Note that before conducting the experiments, the entire cell population was pre-incubated with photoactivatable dyes (which did not affect the TP mitosis ratio; **Figure S3**, **Experimental Section**). The cells of interest became fluorescent upon selective patterned illumination (**Figure 1c**), a feature integrated in our UFO microscopic setup^[13]^. Following this, the entire cell population was detached, and fluorescence-activated cell sorting (FACS) was used to collect fluorescently phototagged CIN cells alongside cells not exhibiting CIN phenotypes. Subsequently, these cells underwent single-cell omics sequencing, including single-cell genomic sequencing (scDNAseq) and transcriptomic sequencing (scRNAseq).

### 2.2 Progeny cells resulting from tripolar mitosis exhibit either polyploidy or high chromosomal variability

We applied CIN-seq to investigate the rare occurrence of spontaneous multipolar mitosis, the most dominant CIN phenotype in MCF10A cells, which typically occurs at a rate of ∼1-2% (**Supporting information and Figure S4**), focusing particularly on the most prevalent form: tripolar mitosis. We observed a high survival rate among TP cells (99.4%, defined as surviving for > 24 hrs or until the next mitosis; **Figure S5**), akin to cells undergoing bipolar mitosis (referred to as BP cells). This contrasts with the high lethality seen in induced multipolar mitosis^[3]^ and aligns with the previously observed high survival rate in cases of spontaneous multipolar mitosis^[8]^, which can further propagate chromosome aneuploidy to their progeny cells. Moreover, the mechanisms underlying tripolar mitosis and the high survival rate of TP cells had remained unclear up to now, prompting further investigation into their biological differences at the molecular level and the elucidation of the underlying mechanisms through our CIN-seq data. Given the sparse nature of TP cells, (384-well) plate-based scDNAseq and scRNAseq were employed in the CIN-seq pipeline (**Experimental Section**). CNV analysis from scDNAseq revealed original, stable, clonal CNVs inherited in MCF10A cells (**Figure 2a**), with higher CNVs in Chromosome (Chr) 8 and partially higher CNVs in Chromosomes (Chrs) 1, 5, 7 and 19 in both BP and TP cells, consistent with previous studies^[15]^. TP cells exhibited two distinct CNV patterns: chaotic and non-chaotic, each accounting for ∼50% of TP cells (**Figure 2a**). Chaotic TP cells had mosaic CNV profiles with euploidy similarity scores below 0.8 (**Experimental Section**), while non-chaotic TP cells had CNV profiles similar to BP cells, with scores approaching 1. This phenomenon resulted in higher heterogeneity and aneuploidy across all chromosomes within TP cells compared to BP cells (**Figure 2b**). To validate the chaotic CNV pattern, a FISH assay using Chromosome 6 (N = 20 cells; **Figure 2c**) revealed similar findings to scDNAseq-derived CNV profiles. Some TP cells retained diploidy or euploidy of Chr 6 (∼36%), while others displayed random aneuploidy (∼64%) (**Figure 2c; Figure S6**). Summing the chaotic CNV profiles from all TP cells bearing chaotic CNVs (referred to as TP-C cells) reconstructed the “normal” CNV pattern observed in BP cells but with an overall higher copy number (∼1.3 times higher) (**Figure 2d**). This supports the existence of chaotic CNV profiles in TP cells. Initially, TP cells with non-chaotic CNV patterns (referred to as TP-NC cells) were thought to be diploid, similar to BP cells (**Figure 2a**). However, when correlating the FACS intensity profiles of those TP cells with the CNV profiles, we found that ∼75% TP-NC cells exhibited polyploidy, predominantly ranging from tri-to octo-ploidy (**Figure 2e, Figure S7**). This polyploidy phenotype suggests the occurrence of cytokinesis failure either during or following tripolar mitosis^[16]^, wherein multiple copies of chromosomes aggregated within single cells, as confirmed by our live-cell imaging (**Figure S8a and S8b**) and FISH assay (N = 20, **Figure S8c**). Collectively, CIN-seq-mediated scDNAseq provides direct evidence of the significant cell-to-cell variability in chromosomal gains and losses, characteristics of CIN. This variability was particularly observed in chaotic TP cells, with non-chaotic TP cells displaying a polyploid karyotype.

**Figure 2.**
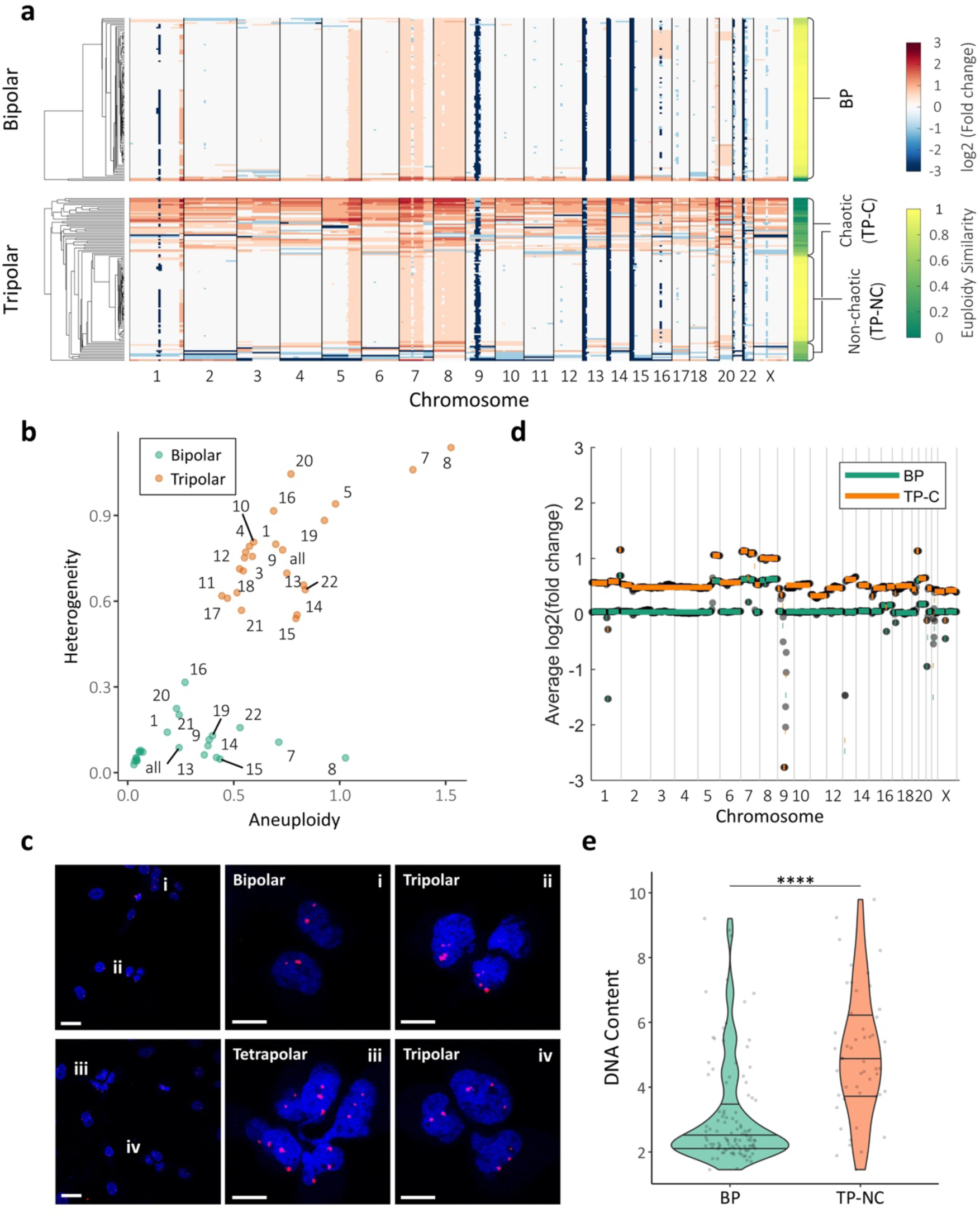
Tripolar mitosis resulted in progeny cells exhibiting either polyploidy or high chromosomal variability. a) CNV profiles, derived from CIN-seq mediated single-cell DNA sequencing, are depicted for cells undergoing bipolar (upper panel) or tripolar mitosis (lower panel). Each row represents an individual cell, while each column illustrates different chromosomes. CNVs are visualized in a red-blue color heatmap, with red indicating higher CNVs in chromosome segments and blue indicating lower CNVs. The euploidy similarity score, displayed on the yellow green color bar, was calculated for each cell. Non-chaotic tripolar dividing cells (**TP-NC**) exhibit a euploidy similarity score close to 1, whereas chaotic tripolar dividing cells (**TP-C**) display a euploidy similarity score less than 0.8. b) A scatter plot illustrates the relationship between the aneuploidy score (x axis) and heterogeneity score (y axis) for CNVs of individual chromosomes in cells undergoing bipolar (green) or tripolar (orange) mitosis. The aneuploidy score measures divergences from euploidy, while the heterogeneity score quantifies the extent of karyotypic heterogeneity. c) Fluorescence in situ hybridization (FISH) assay targeting Chromosome 6 (Chr 6) revealed the chromosome count in different mitosis scenarios: bipolar mitosis (i), tripolar mitosis (ii, iv), and tetrapolar mitosis (iii). Scale bars in the left two graphs denote 30 µm, while those in the inset graphs indicate 10 µm. d) The average of the chaotic CNV profiles from TP cells harboring chaotic CNVs (**TP-C** cells) recapitulates the “normal” CNV pattern observed in control bipolar mitosis (**BP**) cells, with an overall higher copy number (∼1.3 times higher). e) DNA content distributions from FACS intensity of BP cells and TP cells carrying non-chaotic CNVs (**TP-NC** cells). The p-value was obtained using Student’s t-test. ns (not significant); p < 0.05 *; p < 0.01 **. The overlaid quantiles (25th, 50th, 75th percentiles) represent the data spread. Individual data points are shown as jittered points.

### 2.3 The aneuploidy in Chromosome 16 is correlated with a higher rate of tripolar mitosis

Moreover, a subset of BP and TP cells was found to exhibit elevated CNVs of Chrs 16 and 20 (**Figure 2a**). To investigate whether higher CNVs of these chromosomes correlate with multipolar mitosis, we employed CIN-seq to isolate individual TP cells alongside control BP cells. This process was followed by clonal amplification over multiple generations and subsequent bulk DNA sequencing (**Figure 3a**). Clonal expansion originating from TP cells highlighted their ability to generate viable cell populations capable of driving tumor cell proliferation, albeit with a slightly lower proliferation rate (**Figure S9**) but a higher proportion of surviving clones derived from TP cells (**Figure S10**). Furthermore, among the six amplified TP cell clones, none demonstrated elevated CNVs in Chr 20, while only two clones (Clones 1 and 2 – C1 and C2) exhibited higher CNVs in Chr 16 (**Figure 3a**). In contrast to stable chromosome gains (Chrs 1, 5, 7, 8, and 19), these findings suggest that Chrs 16 and 20 gains are inherently unstable and are inefficiently transmitted to all progeny cells. This occurs despite a subset of cells within the amplified TP cell clones carrying higher CNVs in Chr 16. To determine whether the additional gain of Chr 16 observed in Clones C1 and C2 correlates with tripolar mitosis, we quantified the tripolar mitosis rate in these clones compared to the control clones exhibiting bipolar mitosis (Ctrl) (**Figure 3b**). Our analysis revealed that Clones C1 and C2, characterized by elevated CNVs in Chr 16, exhibited a higher rate of tripolar mitosis compared to the control clone. We hypothesized that the aneuploidy of Chr 16 might underlie this correlation. To validate this hypothesis, we employed CRISPR imaging technology^[17]^ to visualize Chr 16 counts during live bipolar and multipolar mitoses. In this assay, catalytically inactive dCas9 was fused to three mCherry copies (dCas9-3XmCherry) and stably expressed in MCF10A cells (**Experimental Section**). Introducing single-guide RNA sequences targeting the centromeric sequence of Chr 16^[18]^ (TGGATATCTTGGCCTCTTAG) enabled dCas9-3XmCherry to aggregate at the Chr 16 centromere. By quantifying aggregated fluorescent spots or foci prior to and during mitosis, we determined the number of Chr 16 copies involved in tripolar and bipolar mitoses. The analysis revealed a statistically significant increase in the number of Chr 16 copies, particularly uneven numbers, in TP mother cells compared to bipolar mother cells (**Figure 3c and 3d**). As an additional validation step, we quantified the number of Chr 6 counts, a diploid chromosome in MCF10A cells, and confirmed that Chr 16 indeed exhibited significantly higher chromosome counts per nucleus compared to Chr 6 (**Figure 3e and 3f**). These results confirm that aneuploidy in Chr 16 is strongly associated with a higher tripolar mitosis rate, thereby validating our initial hypothesis.

**Figure 3.**
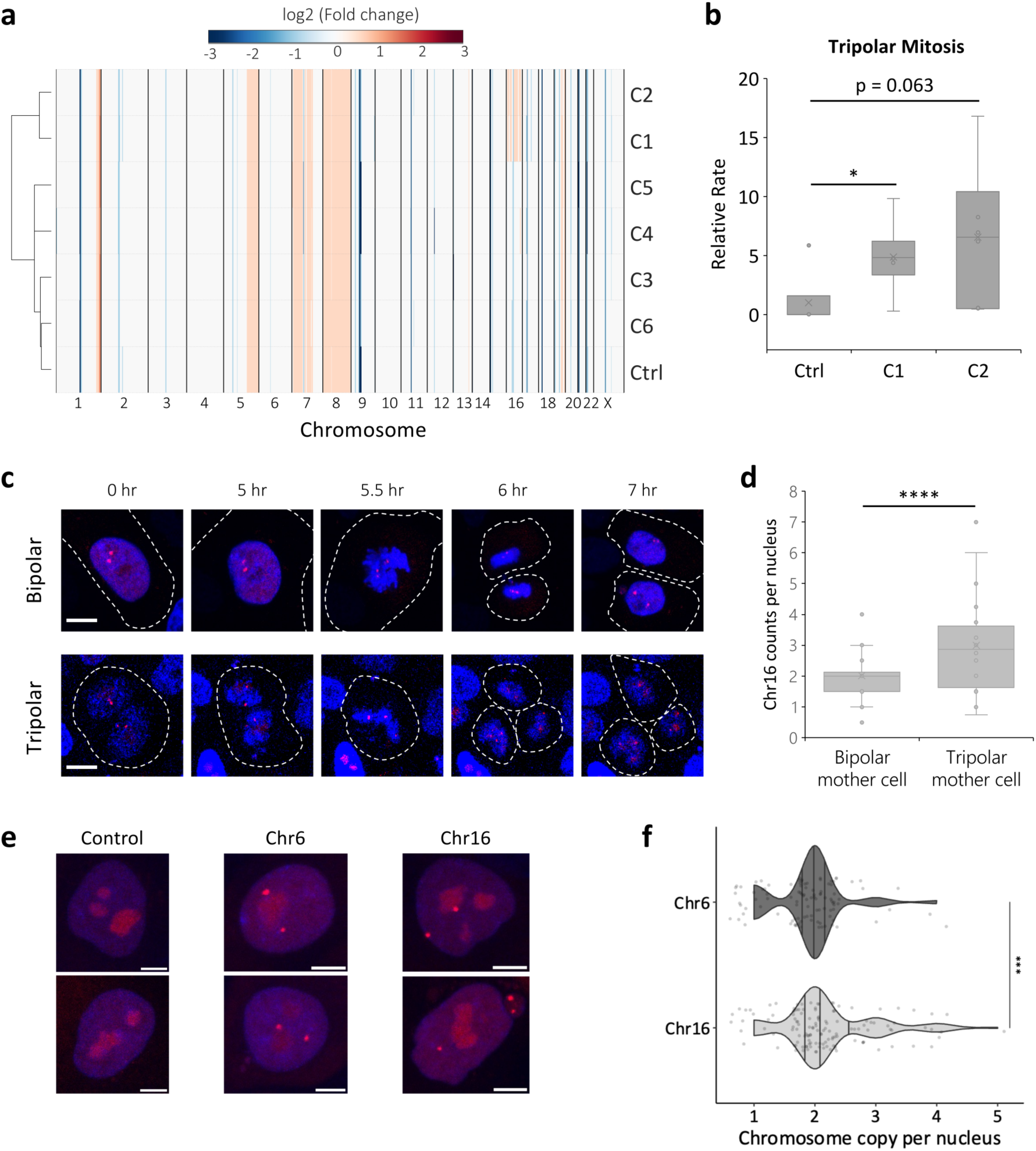
Aneuploidy in Chromosome 16 is linearly correlated with an increased tripolar mitosis rate. a) CNV analysis of bulk DNA sequencing, derived from 6 amplified TP cell progenies (C1-C6) collected via CIN-seq, as well as a control (Ctrl) clone (randomly isolated cell clone). CNVs are visualized in a red-blue color heatmap, with red indicating higher CNVs in chromosome segments and blue indicating lower CNVs. b) Box and whisker plot of tripolar mitosis rates for clones C1, C2, and Ctrl corresponding to the cell clones shown in Figure 3e. The box shows the interquartile range (25th, 50th, 75th percentiles), and whiskers indicate the minimum and maximum values. The p-value was obtained using Student’s t-test. ns (not significant); p < 0.05 *; p < 0.01 **; p < 0.001 ***; p < 0.0001 ****. c) CRISPR imaging of MCF10A cells expressing dCas9-3XmCherry, which forms fluorescent foci (red spots) upon the addition of single-guide RNAs (sgRNAs) targeting the centromeric sequence of Chr 16. Representative cells undergoing bipolar (top 5 panels) or tripolar (bottom 5 panels) mitosis are shown. White dashed lines indicate cell membranes. Scale bars denote 10 µm. d) Box and whisker plot of Chromosome 16 counts, measured from CRISPR imaging (N = 20 cells for TP cells, and N = 30 cells for BP cells) for mother cells undergoing bipolar or tripolar mitosis. The box shows the interquartile range (25th, 50th, 75th percentiles), and whiskers indicate the minimum and maximum values. The p-value was obtained using Student’s t-test. ns (not significant); p < 0.05 *; p < 0.01 **; p < 0.001 ***; p < 0.0001 ****. e) CRISPR imaging of MCF10A cells expressing dCas9-3XmCherry, which forms fluorescent foci (red spots) upon the addition of sgRNAs targeting the centromeric sequence of Chr 6 or 16. Representative cells without the addition of sgRNAs (left 2 panels), with sgRNAs targeting Chr 6 (middle 2 panels), or with sgRNAs targeting Chr 16 (right 2 panels) are shown. Scale bars denote 10 µm. f) CRISPR imaging assay to quantify the average number of Chr 6 or Chr 16 counts per nucleus (N = 103 for Chr 6 and N = 132 for Chr 16). The p-value was obtained using Student’s t-test. ns (not significant); p < 0.05 *; p < 0.01 **; p < 0.001 ***; p < 0.0001 ****. Statistical analysis was carried out using R Software and Microsoft Excel.

### 2.4 RhoGTPase Cycle signaling pathways drove cytokinesis failure, resulting in the formation of polyploid cells

To gain insight into the underlying molecular mechanisms of tripolar mitosis and explore the differences between TP-C and TP-NC cells, we employed CIN-seq to collect BP cells and TP cells separately for scRNAseq (**Experimental Section**). Through Seurat clustering analysis, we identified 3 distinct clusters on UMAP embedding (**Figure 4a**). Mapping these clusters to the phenotype data revealed that Seurat cluster 0 (S0) predominantly comprised BP cells, Seurat cluster 1 (S1) consisted of a mix of both BP and TP cells, and Seurat cluster 2 (S2) was predominantly composed of TP cells (**Figure 4a**). We hypothesized that the two TP cell subtypes observed in S1 and S2 clusters might be related to TP cells displaying two distinct CNV profiles (TP-C or TP-NC cells) observed in the scDNAseq data. By correlating their FACS intensity profiles, we found that TP cells in the S1 cluster (TP-S1) closely resembled TP cells with a non-chaotic CNV pattern (TP-NC cells), whereas TP cells in the S2 cluster (TP-S2) exhibited similarities to TP cells with a chaotic CNV pattern (TP-C cells) (**Figure 4b**). We have shown that the polyploidy observed in TP-NC cells (a.k.a. TP-S1 cells) resulted from failed cytokinesis (**Figure 2e; Figure S7**). The BP cells grouped within the same S1 cluster (BP-S1 cells, **Figure 4c**) as TP-S1 cells were also likely polyploid, undergoing cytokinesis failure similar to TP-S1 cells. To investigate the mechanisms underlying cytokinesis failure, we conducted differential gene expression and pathway analyses comparing polyploid TP-S1 and BP-S1 cells with diploid BP cells (bipolar cells from the S0 cluster, BP-S0) (**Figure 4d**). This analysis revealed a significant upregulation of the RhoGTPase cycle pathway in both polyploid BP-S1 and TP-S1 cells (**Figure 4d and 4e**). While the observed upregulation of Rho GTPase cycle signaling in polyploid cells could represent either a driving mechanism or a downstream consequence, the significant increase in cytokinesis failure upon activation of the Rho GTP signaling pathway using the Rho/Rac/Cdc42 activator (**Figure 4f, Figure S11, Experimental Section**) suggests that the upregulated Rho GTP signaling is associated with cytokinesis failure. Rho GTPase cycle signaling regulates actin and cytoskeletal dynamics^[19]^, which was also found to be upregulated in polyploid BP-S1 and TP-S1 cells (**Figure 4d**). Dysfunction in these pathways may contribute to the failed cytokinesis observed in our data. Interestingly, despite the activation of Rho GTP signaling, there was no corresponding increase in the incidence of tripolar mitosis (**Figure 4g**). This finding indicates that while Rho GTP signaling mediated cytokinesis failure contributes to polyploidy, it is insufficient on its own to induce tripolar mitosis.

**Figure 4.**
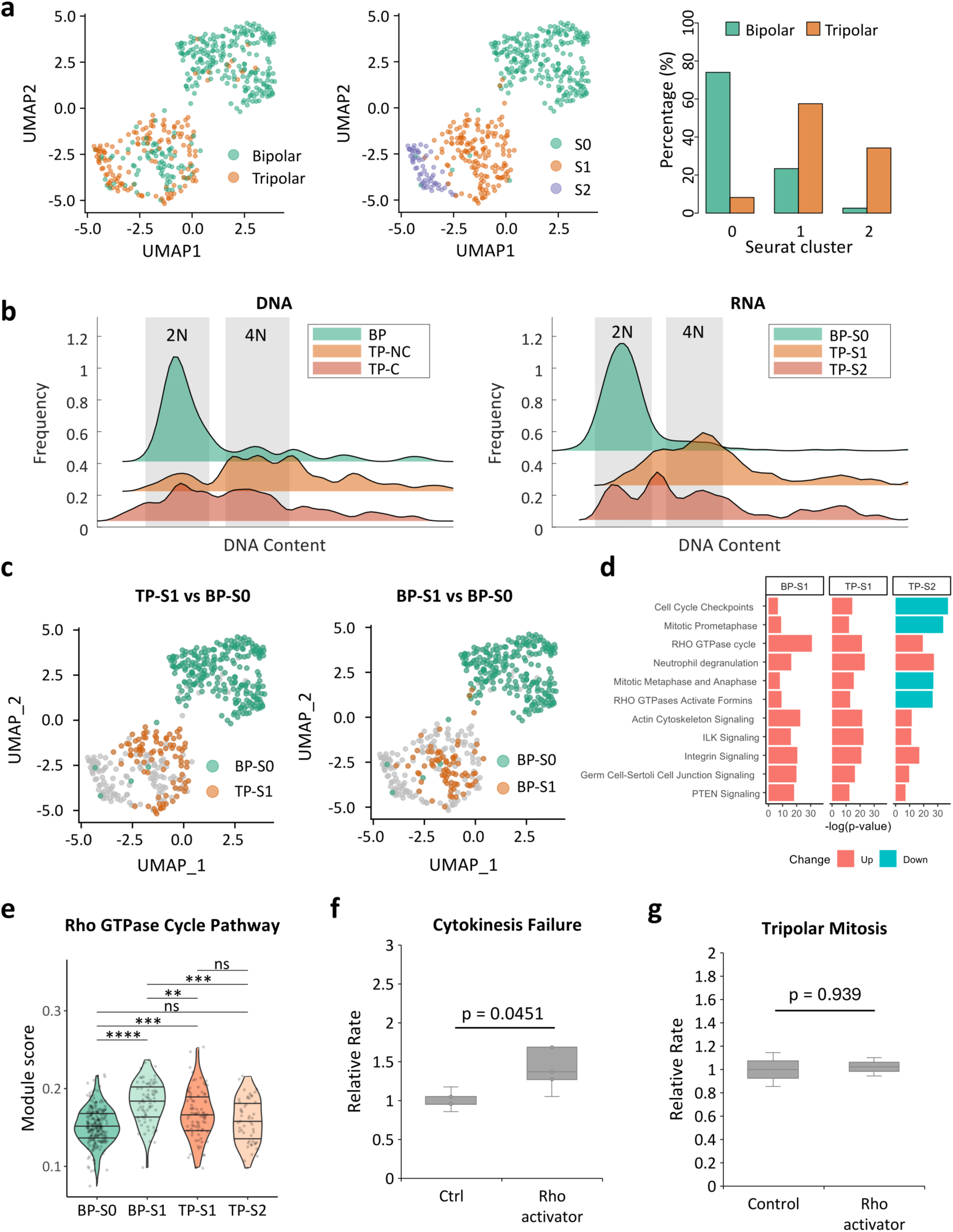
RhoGTPase signaling drove cytokinesis failure, resulting in the formation of polyploid cells. a) UMAP plots illustrate cells exhibiting either mitosis phenotypes (left panel: bipolar mitosis in green or tripolar mitosis in orange) or Seurat-based clusters (middle panel: Cluster 0-2, S0-S2). Right panel: a bar plot showing the composition (in percentage) of bipolar or tripolar mitosis phenotype within each Seurat cluster. b) Flow cytometry plots show the correlation between chaotic and non-chaotic CNV cells and TP cells in Seurat clusters 1 and 2 based on their DNA content. c) UMAP plots display the results of the differential gene analysis conducted on TP-S1 and BP-S1 cells compared to BP-S0 cells. d) IPA pathway analysis highlights the shared top-ranked pathways identified using DEGs from BP-S1, TP-S1 and TP-S2 cells in comparison to BP-S0 cells. e) A violin plot presents the module score of the RhoGTPase Cycle pathway for each cell subcluster. The p-value was obtained using Wilcoxon test with Bonferroni correction. ns (not significant); p < 0.05 *; p < 0.01 **. The overlaid quantiles (25th, 50th, 75th percentiles) represent the data spread. Individual data points are shown as jittered points. BP-S0: BP cells in Seurat cluster 0. BP-S1: BP cells in Seurat cluster 1. TP-S1: TP cells in Seurat cluster 1. TP-S2: TP cells in Seurat cluster 2. f) Box and whisker plot of cytokinesis failure rates in cells treated with a Rho GTP signaling activator (Control, Ctrl) (N = 3). The box shows the interquartile range (25th, 50th, 75th percentiles), and whiskers indicate the minimum and maximum values. The p-value was obtained using Student’s t-test. ns (not significant); p < 0.05 *; p < 0.01 **; p < 0.001 ***; p < 0.0001 ****. g) Box and whisker plot of tripolar mitosis rates in cells treated with a Rho GTP signaling activator or without (Control, Ctrl) (N = 3) The p-value was obtained using Student’s t-test. The box shows the interquartile range (25th, 50th, 75th percentiles), and whiskers indicate the minimum and maximum values. The p-value was obtained using Student’s t-test. ns (not significant); p < 0.05 *; p < 0.01 **; p < 0.001 ***; p < 0.0001 ****. Statistical analysis was carried out using R Software and Microsoft Excel.

### 2.5 Degranulation-like signaling is activated in TP cells and associated with a stress-responsive lysosomal program

Through CIN-seq-mediated single-cell transcriptomic analysis, we identified a significant activation of neutrophil degranulation-like signaling in TP cells, particularly in the chaotic TP-S2 (TP-C) subtype (**Figure 5a, b**). Although degranulation is classically a hallmark of innate immune cells such as neutrophils^[20]^, no immune cells were present in our assays, suggesting that cancer cells themselves can activate a similar secretory response under mitotic stress. In neutrophils, degranulation involves the release of granule-stored enzymes and proteases into the extracellular milieu to degrade pathogens and damaged material^[20]^. Analogously, we hypothesize that TP cells, experiencing extreme chromosomal instability and accumulation of malfunctioning or misfolded proteins following aberrant cytokinesis, invoke a comparable intracellular degranulation-like mechanism to manage proteotoxic stress. Supporting this idea, several enzymes or related proteins typically associated with granule or lysosomal compartments were among the most significantly upregulated genes in TP versus BP cells (**Figure 5a**), including CTSD (*cathepsin D*), CSTB (*cystatin B*), CDA (*cytidine deaminase*), and PYGB (*glycogen phosphorylase B*) (highlighted in red in **Figure S12**). Functional annotation of these genes revealed enrichment in degranulation and lysosomal secretion pathways, suggesting activation of a stress-induced lysosomal proteolytic network following multipolar mitosis.

**Figure 5.**
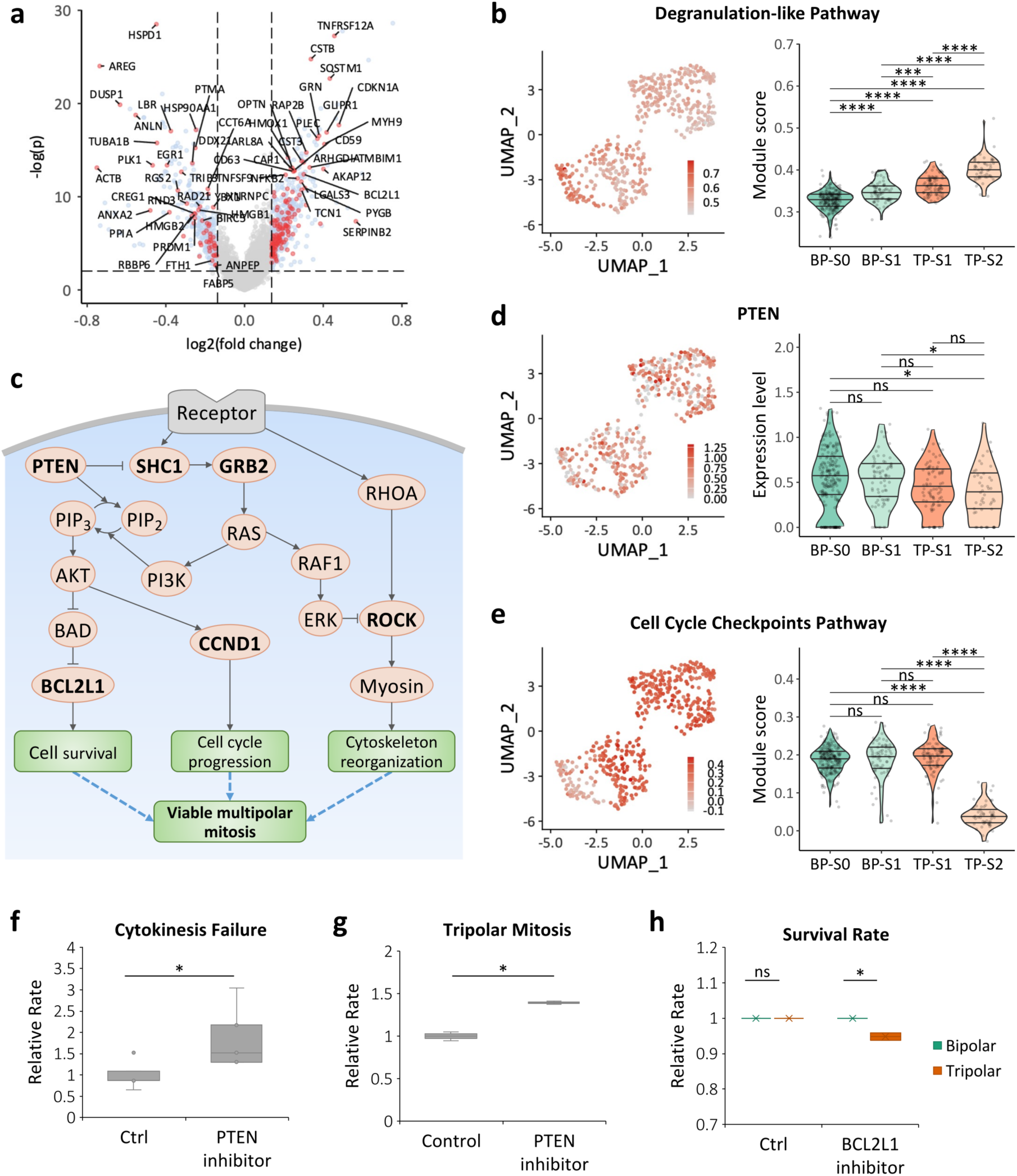
Transcriptomic analysis derived from CIN-seq (tripolar mitosis). a) A volcano plot displays the results of the differential gene analysis conducted on TP cells compared to BP cells. Genes associated with the degranulation-like signaling and PTEN signaling pathways were highlighted. b) A UMAP plot displays the module score of the degranulation-like pathway (left panel). A violin plot presents the module score of this pathway for each cell subcluster (right panel). The p-value was obtained using Wilcoxon test with Bonferroni correction. ns (not significant); p < 0.05 *; p < 0.01 **. The overlaid quantiles (25th, 50th, 75th percentiles) represent the data spread. Individual data points are shown as jittered points. BP-S0: BP cells in Seurat cluster 0. BP-S1: BP cells in Seurat cluster 1. TP-S1: TP cells in Seurat cluster 1. TP-S2: TP cells in Seurat cluster 2. c) A mechanistic schematic elucidates the role of PTEN-associated signaling pathway in tripolar mitosis. d) A UMAP plot (left panel) alongside a violin plot (right panel), illustrating the expression of the PTEN gene across various cell subclusters. The p-value was obtained using Wilcoxon test with Bonferroni correction. ns (not significant); p < 0.05 *; p < 0.01 **. The overlaid quantiles (25th, 50th, 75th percentiles) represent the data spread. Individual data points are shown as jittered points. e) A UMAP plot illustrates the module score of the Cell Cycle Checkpoints pathway (left panel). A violin plot depicts the module score of this pathway for each cell subcluster (right panel). The p-value was obtained using Wilcoxon test with Bonferroni correction. ns (not significant); p < 0.05 *; p < 0.01 **. The overlaid quantiles (25th, 50th, 75th percentiles) represent the data spread. Individual data points are shown as jittered points. f) Box and whisker plot of cytokinesis failure rates in cells treated with a PTEN inhibitor or without (Control, Ctrl) (N = 3). The box shows the interquartile range (25th, 50th, 75th percentiles), and whiskers indicate the minimum and maximum values. The p-value was obtained using Student’s t-test. ns (not significant); p < 0.05 *; p < 0.01 **; p < 0.001 ***; p < 0.0001 ****. g) Box and whisker plot of tripolar mitosis rates in cells treated with a PTEN inhibitor or without (Control, Ctrl) (N = 3). The box shows the interquartile range (25th, 50th, 75th percentiles), and whiskers indicate the minimum and maximum values. The p-value was obtained using Student’s t-test. ns (not significant); p < 0.05 *; p < 0.01 **; p < 0.001 ***; p < 0.0001 ****. h) Box and whisker plot of survival rates for BP (green) and TP (orange) cells, comparing treatment outcomes with or without a BCL2L1 inhibitor (N = 3). The box shows the interquartile range (25th, 50th, 75th percentiles), and whiskers indicate the minimum and maximum values. The p-value was obtained using Student’s t-test. ns (not significant); p < 0.05 *; p < 0.01 **; p < 0.001 ***; p < 0.0001. Statistical analysis was carried out using R Software and Microsoft Excel.

To functionally evaluate the role of this response, we inhibited the lysosomal aspartyl protease CTSD using Pepstatin (**Figure S13**). Based on the proteostasis hypothesis, we initially expected that CTSD inhibition would compromise clearance of damaged peptides and thus reduce cell viability. Surprisingly, however, Pepstatin treatment led to a significant increase in the survival of TP-derived daughter cells compared to untreated controls. This unexpected outcome indicates that CTSD activity does not act solely as a cytoprotective degradation mechanism but may also contribute to a pro-death lysosomal stress pathway triggered by severe genomic instability. Together, these findings suggest that the coordinated upregulation of degranulation-associated enzymes in TP cells represents a stress-activated lysosomal program that functions as a cellular quality-control checkpoint to eliminate severely damaged progeny. Inhibition of this pathway, as observed with CTSD blockade, permits transient escape from lysosomal cell death, thereby enhancing survival following multipolar mitosis.

### 2.6 PTEN attenuation and BCL2L1 upregulation cooperatively promote survival of TP cells despite lysosomal stress

Integrative network and pathway analyses further revealed that TP cells exhibited extensive transcriptional remodeling within the PTEN-dependent signaling network (**Figure 5a, Figure S14**), including the upregulated genes SHC1, GRB2, CCND1, and BCL2L1 (**Figure 5a and S14**), and the downregulated genes PTEN, ROCK1, and ROCK2 (**Figure 5a, Figure S14**). Building on these observations and the findings from the network analysis, we hypothesized the mechanisms driving tripolar mitosis and their contributions to the elevated survival rates, as outlined in the proposed mechanism depicted in **Figure 5c**. Specifically, the downregulation of PTEN (**Figure 5c and 5d**) led to the upregulation of the SHC1 and GRB2 genes (**Figure 5c and S14**), which indirectly elevated the expression of CCND1 (**Figure 5c and S14**), a gene pivotal for cell cycle progression^[21]^ (**Figure 5c**). Additionally, we observed a unique downregulation of the cell cycle checkpoints pathway (**Figure 4d and 5e**), particularly evident in TP-S2 (a.k.a. TP-C) cells, which harbored chaotic CNV profiles. The cell cycle checkpoints pathways play a crucial role in preventing cells from entering mitosis in the presence of mis-segregated chromosomes or failed cytokinesis, mechanisms vital for safeguarding genome integrity^[22]^. This finding, along with the upregulated CCND1 gene (**Figure 5c and S14**), provides insights into the successful and continued completion of the mitosis process in TP cells despite their genomic abnormalities.

Additionally, according to the network analysis, the upregulation of SHC1 and GRB2 following PTEN downregulation was indirectly linked to the inhibition of the ROCK genes (ROCK1 and ROCK2) (**Figure 5c and S14**), which play a critical role in cytoskeletal regulation^[23]^. This inhibition has been shown to disrupt cytoskeletal reorganization^[24]^, potentially resulting in failed cytokinesis and subsequent multipolar mitosis, as observed in a subset of TP cells (TP-S1). We validated this by observing elevated occurrences of both cytokinesis failure and tripolar mitosis upon inhibiting the PTEN gene (**Figure 5f, g, Figure S15, Experimental Section**). Among the genes upregulated within this network, *BCL2L1*, encoding the anti-apoptotic protein BCL-xL^[25]^, was significantly upregulated in TP cells (**Figure 5c, Figure S14**), and likely plays a pivotal role in their high survival rate (99.4%; **Figure S5**). We confirmed the essential role of *BCL2L1* in TP-cell viability by demonstrating that pharmacological inhibition of BCL2L1 reduced survival specifically in TP cells, but not in BP controls (**Figure 5h, Figure S16**).

To further validate the relationship between PTEN and BCL2L1, we performed the following functional assay. We compared the survival rates of TP cells in MCF10A cells with PTEN-wild type (WT), PTEN-overexpression (OE), or PTEN-OE combined with BCL2L1-OE. The results showed that PTEN overexpression significantly reduced the survival rate of TP cells compared with the PTEN-WT group (**Figure S17**). Co-overexpression of BCL2L1 rescued the decreased survival caused by PTEN overexpression, and no significant difference was observed between the PTEN-WT and PTEN-OE/BCL2L1-OE groups. These results indicate that BCL2L1 acts downstream of PTEN to promote the survival of tripolar mitosis cells, and that loss of PTEN enhances cell viability primarily through BCL2L1-mediated anti-apoptotic signaling, thereby establishing a functional link between PTEN attenuation and the survival advantage of multipolar mitosis progeny.

Moreover, given the concurrent activation of CTSD-associated lysosomal stress programs, increased BCL-xL expression likely acts as a protective buffer, restraining CTSD-mediated apoptotic execution and enabling TP-derived cells to survive despite activation of pro-death lysosomal signals. This interpretation is consistent with our functional findings, where CTSD inhibition further enhanced TP-cell survival (**Figure S13**), suggesting that lysosomal stress and anti-apoptotic signaling coexist in a finely balanced state.

Furthermore, when BCL2L1 was pharmacologically inhibited under CTSD-inhibited conditions, the previously enhanced TP-cell survival was reversed, restoring cell viability to baseline levels (**Figure S18**). This reversal confirms that BCL2L1 activity is essential for sustaining TP-cell survival even in the context of CTSD-associated lysosomal stress, highlighting its pivotal role in buffering apoptotic pressure in TP cells. Collectively, these results support a unified model (**Figure 5c**) in which PTEN downregulation is closely associated with viable and successful tripolar mitosis, facilitated by cell cycle progression, suppression of cell cycle checkpoint pathways, and enhanced cell survival mediated by BCL2L1 (**Figure 5c**).

To assess whether the findings from MCF10A cells, namely that upregulation of RhoGTPase signaling and downregulation of PTEN contribute to cytokinesis failure and tripolar mitosis, could also be observed in a cancerous cell line, we conducted similar assays using MCF7 breast cancer cells. Activation of the RhoGTPase pathway in MCF7 cells resulted in a higher incidence of failed cytokinesis under the treatment conditions (**Figure S19a**; **Experimental Section**). Additionally, PTEN attenuation led to an increased occurrence of failed cytokinesis and tripolar mitosis (**Figure S19**), mirroring the results observed in MCF10A cells. These outcomes further confirm the validity of the CIN-seq discoveries and elucidate the underlying mechanisms driving cytokinesis failure and tripolar mitosis.

## 3. Conclusion

In summary, multipolar mitosis, a prominent manifestation of chromosomal instability, often arises spontaneously in cancers, comprising rare subpopulations of cells alongside other CIN phenotypes like lagging chromosomes and chromosomal bridges. However, systematically profiling these rare and dynamic cell subsets has been a significant challenge. In this study, we introduced CIN-seq, a novel approach that enables high-throughput single-cell genomic and transcriptomic sequencing of these rare, dynamic CIN subpopulations on a larger scale for the first time. This advancement is made possible by the implementation of larger field-of-view UFO microscopy, real-time GTWeka cell segmentation, instant detection of single cells displaying abnormal mitotic phenotypes, and automated phototagging of such cells during time-lapse recordings. As CIN-seq enables the direct linking of CIN phenotypes with sequenced genomic and transcriptomic profiles, it unveils novel insights that are typically obscured by standard single-cell sequencing methods. These traditional methods often lack the capability to link phenotypes to genotypes, let alone to rare cell subpopulations exhibiting live, dynamic CIN phenotypes observed during the mitotic phase. Here, we applied CIN-seq to profile MCF10A exhibiting live, spontaneous tripolar mitosis in action, revealing several novel insights into the underlying mechanisms of this phenomenon. Our CIN-seq approach represents a groundbreaking advancement in CIN research, providing a powerful tool to capture CIN-associated temporal RNA expression or newly generated RNA transcripts. This approach not only provides a new methodology to address pressing clinical challenges, but also offers novel insights unattainable with traditional techniques. The impact of the technology can be extended to decipher unique molecular mechanisms tailored to other specific CIN phenotypes (e.g., lagging chromosomes) or tumor types, particularly in patient derived tumor samples, and to pinpoint the most relevant signaling pathways behind such events. Through this approach, we unlock novel insights into the mechanisms underlying naturally occurring chromosomal instability, thereby paving the way for the identification of new therapeutic strategies. Historically, therapeutic interventions targeting CIN have faced limited progress^[3]^, largely due to the incomplete understanding of this phenotype. However, with the comprehensive insights provided by CIN-seq, we have the potential to accelerate advancements in the field of cancer therapeutics.

## 4. Experimental Section

### 4.1 Cell Lines and Cell cultures

#### MCF10A cells

MCF10A cells (CVCL_0598), a gift from Dr. Reuven Agami (Dutch National Cancer Institute, NKI), were cultured in DMEM/F12 medium (ThermoFisher, 11320033) supplemented with 5% horse serum (Fisher Scientific, SH3007403), 1% Penicillin/Streptomycin, 1% (*w:w*) insulin (ThermoFisher, 12585014), 0.05% (*w:w*) hydrocortisone (Stem Cell, 74142), 0.01% (*w:w*) cholera toxin (Sigma, SAE0069) and 0.001% epidermal growth factor (ThermoFisher, PHG0313). The cultures were maintained in a 37 °C incubator under 5% CO2. The cell line was free of contamination, as it was routinely tested, including for mycoplasma.

#### MCF7 cells

MCF7 cells were cultured in DMEM, Dulbecco’s Modified Eagle Medium (ThermoFisher, 21063) supplemented with 10% Fetal Bovine Serum (Capricorn, FBS-12A) and 1% Penicillin/Streptomycin. The cultures were maintained in a 37 °C incubator under 5% CO2.

### 4.2 Chromosome Instability Sequencing (CIN-seq)

Pipeline overview. Two to five days (cell-dependent) before conducting experiments, cells were seeded on culture dishes in culture medium, and the medium was supplemented with 1X SPY650-DNA nuclear stain (Spirochrome, SC501) the night before the assay. On the day of the assay, the culture medium was further supplemented with 15-30 μM photoactivatable dyes, PA Janelia Fluor 549 (Tocris, 6147), for 10-15 min. Subsequently, the cells were washed three times with pre warmed PBS before being incubated with fresh culture medium and placed in the cell incubator of the ultrawide field-of-view optical (UFO) microscope [12] for live-cell, multi-region, time-lapse recording for 2-5 hrs (in this study) or longer (depending on the phenotype of interest).

The UFO microscope is a custom-built imaging system designed for high-throughput screening of tens of thousands of cells per field-of-view with subcellular resolution [12]. Our speedy mitosis detection algorithm (see the ‘Real-time CIN detection’ section) was executed in real-time for every two frames of a region-of-interest, and the coordinates of target cells (cells displaying tripolar mitosis or micronuclei) were recorded. The photoactivatable dyes in the target cells were photoactivated via a 405 nm laser for 0.2 seconds, automatically steered by galvanometer scanners at UFO, which received the target cell coordinates from real-time image analysis. Once all target cells were photoactivated, they were detached using 1X trypsin-EDTA and collected in Hanks’ balanced salt solution (HBSS). The cells were spun down to remove excess trypsin and resuspended in HBSS at a cell concentration of approximately 1000 cell/μL. Subsequently, the BD FACSMelody cell sorter was employed to sort cells using yellow/green (561 nm) and red lasers (640 nm), with filters for PE (582/15 nm) and APC (660/10 nm), respectively. Cells exhibiting positive signals in the PE channel, indicative of photoactivation, were sorted into a microwell plate for subsequent single-cell sequencing.

### 4.3 MCF10A cell preparation for the FUNseq assay

Two days before conducting experiments, approximately 100,000 cells were seeded on a fibronectin-coated 6-well glass-bottom plate (Cellvis) in culture medium supplemented with 1X SPY650-DNA nuclear stain (Spirochrome). Following an overnight incubation to allow cell attachment, the medium was supplemented with 9 μM RO-3306 (Sigma Aldrich), a compound known for its ability to synchronize the cell cycle before the G2/M phase. After 17 hr, the culture medium was supplemented with 15 μM photoactivatable dyes for 15 min. Subsequently, and the cells were washed three times with pre-warmed PBS before being incubated with new culture medium without RO-3306. Upon release from cell cycle synchronization, the MCF10A cells were placed in the cell incubator of the UFO microscope for the FUNseq assay.

### 4.4 Real-time mitosis detection

The raw images underwent initial processing for cell segmentation using our previously developed ground truth-assisted trainable Weka segmentation (GTWeka) [13]. In this process, three optimal image filters, namely 2-σ Gaussian blur, 4-σ Sobel filter and 2-σ eigenvalues of Hessian matrix, were applied to the raw image to generate three filtered images, or image features. Subsequently, for each individual pixel (1.22 μm/pixel) in the raw image, three newly calculated values were obtained from the image features. Using combinations of these three values, random forest classification was performed to classify each pixel as either cell or background. This classification yielded a binary cell segmentation map for all pixels in the raw image with all frames in a movie being cell segmented. Following cell segmentation, cells displaying tripolar mitosis were identified using the fixed-radius near neighbor search: for each time point t, every cell was analyzed to identify its own self or daughter cells in the subsequent time point t + 1 within 7 pixels radius (D_MD_). In cases where a cell had more than one cell present in time point t + 1, the distances between daughter cells were examined to determine if they were shorter than 12 pixels (D_DD_). These pixel values were determined from average distance values obtained from processing a large population of 10^5-6^ cells. Daughter cell counts of two were labeled as bipolar mitosis, while counts exhibiting three were classified as tripolar mitosis. The positions of the mother cell, daughter cells and corresponding time points were all recorded throughout the analysis process. The script will be available on GitHub before publication.

### 4.5 CIN-seq-mediated single-cell DNA sequencing and analysis

For single-cell DNA sequencing, cells processed through the CIN-seq pipeline were sorted via FACS into 384-well plates pre-filled with mineral oil. In total, 456 BP cells and 235 TP cells were collected (4 batches). The plates were subsequently frozen and processed following the single-cell NlaIII sequencing procedures [1]. In brief, the nucleus in each cell underwent NlaIII restriction enzyme digestion, resulting in DNA fragments that were ligated to barcoded adapters. These ligated fragments were then subjected to in vitro transcription and prepared as sequencing libraries, followed by 150 base pair (bp) paired-end sequencing using the Illumina NextSeq 2000 platform at a mean coverage rate of ∼1-2% per cell (∼60,000 Megabase pair (Mb)/plate). The raw fastq files generated from the sequencing were aligned against the human reference genome (Ensembl release 97) to produce bam files. These bam files were then imported into the AneuFinder R package (v.1.26.0) [25] for copy number variation (CNV) analysis. Default parameters were employed, including the Edivisive method with a fixed bin size of 5 Mb, with further filtering to exclude artifacts caused by high-repeat regions and to correct GC content for CNV calling. After this, cells with low-quality were filtered out based on a spikiness score < 0.4 or a bhattacharyya score > 0.8. After quality control (QC), 243 BP cells and 107 TP cells were used for downstream analyses in this study. CNV profiles were visualized as clustergrams using Matlab. The euploidy similarity score was calculated by the Hamming distance between the bipolar reference CNV profile, which was the median of all bipolar CNV profiles, and individual CNV profiles. Aneuploidy and heterogeneity scores for individual chromosomes, as well as for all chromosomes, were calculated using the AneuFinder package (v.1.26.0). The aneuploidy score represents the degree of divergence from euploidy, while the heterogeneity score reflects the karyotypic variability within the same type of cells.

### 4.6 CIN-seq mediated single-cell RNA sequencing and analysis

For single-cell RNA sequencing, cells processed through the CIN-seq pipeline were sorted via FACS into 384-well plates pre-filled with barcoded primers. In total, 384 BP cells and 219 TP cells were collected (2 batches). These plates were subsequently frozen followed by single-cell RNA library preparation and sequencing (150,000 reads/cell) using the SORT-seq procedures [26]. In brief, mRNA from each cell was reverse transcribed into complementary double-stranded DNA (cDNA). Subsequently, all contents from each well were pooled, followed by in vitro transcription, library preparation, and Illumina sequencing. Raw fastq files obtained from the sequencing were aligned against the reference genome (Homo sapiens, Hg 38) using the Spliced Transcripts Alignment to a Reference (STAR) aligner (v2.7.10). Subsequently, HTseq (v2.0.0) was employed to generate count tables (cell x gene) with expression values. These data underwent processing using the Seurat package (v4.4.0) [27] in R. Quality control measures were applied to exclude low-quality cells, identified by mitochondria ratios surpassing 30%, as well as potential doublets characterized by gene counts exceeding 10,000. After QC, 308 BP cells and 146 TP cells as well as 563 micronuclei cells and 514 non-micronuclei cells were used for downstream analyses in this study. Standard Seurat procedures were employed, including log-normalization, scaling, and the selection of the 2000 most variable genes for dimensionality reduction analyses (e.g., PCA and UMAP plots). Differential gene expression analysis was conducted using the Linear Models for Microarray Data (limma) package (v.3.54.2) [28], calculated using the **10**Benjamini-Hochberg (BH) procedure with a cutoff of |FoldChange| > 1.2 and an adjusted p value < 0.05. Module scores were calculated using the ‘AddModuleScore’ function in the Seurat package. Pathway-correlated genes were binned based on their averaged expression and subtracted from the aggregated expression of randomly selected genes from each bin. Wilcoxon signed-rank tests, adjusted for multiple comparisons using Bonferroni correction, were conducted within each group to calculate the significance of module scores and gene expression levels. Significance was defined as *p* < 0.05. Statistical analysis was carried out using R Software.

### 4.7 CIRSPR imaging

Production of the dCas9-3XmCherry lentivirus. dCas9-3XmCherry lentivirus was generated through the following protocol: ∼ 5 million human embryonic kidney (HEK) 293T cells, cultured in DMEM (ThermoFisher, 21063029) supplemented with 10% Fetal Bovine Serum and 1% Penicillin/Streptomycin, were seeded in a 10 cm culture dish a day before transfection. The culture medium was refreshed right prior to transfection. Transfection was performed according to the Lipofectamine TM 3000 transfection protocol (ThermoFisher #L3000015). Specifically, the cells were transfected with a DNA mixture comprising 8.5 µg pHAGE-TO-dCas9-3XmCherry (Addgene #64108), 4.5 µg psPAX2 (Addgene #12260) and 4.5 µg pMD2.G (Addgene #12259), using Lipofectamine TM 3000. The virus was harvested at 24-hr intervals twice post-transfection, followed by filtration of the medium through a 0.45 μm pore filter. The collected lentivirus was further concentrated through ultra-centrifugation for 2 hr at 20,000 RPM at 4 °C, after which it was stored at −80°C for preservation. Lentiviral transduction and generation of MCF10A-dCas9-3XmCherry cell line. ∼100,000 MCF10A cells were seeded onto a 35 mm cell culture dish a day before transduction. Following this, the culture medium was replaced with pre-warmed medium containing 10 µg/mL polybrene (Millipore, TR-1003-G), after which dCas9-3XmCherry lentivirus containing ∼1 x 10 5 lentiviral particles was added. After a 24-hr period, the medium was refreshed, and the cells were subsequently cultured in a 10 cm culture dish until reaching confluency. Afterward, cells expressing mCherry were selectively collected via FACS. CRISPR imaging experiments. ∼25,000 cells were seeded in each well of an ibidi µ-Slide 8 well high ibiTreat chambered coverslip (ibidi, 80806) a day before performing the experiments. The single-guide RNA (sgRNA) 16-1 [17] (TGGATATCTTGGCCTCTTAG; manufactured by GenScript), targeting α-satellite centromeric repeats of chromosome 16, was transfected according to the Lipofectamine® RNAiMAX (ThermoFisher, 13778030) transfection protocol. Specifically, 15.8 pmol of sgRNA 16-1, diluted in Opti-MEM TM (ThermoFisher, 51985026), was added into the cells. After 48 hr, the medium was refreshed, and a fresh batch of sgRNA dilution was added to the chambers before the time-lapse recording. Time-lapse movies were recorded using Leica confocal microscopy (Leica SP8 AOBS with a DMi8 system). The nuclear signal, represented by H2B-GFP expression in MCF10A cells, was observed by exciting with a 488 nm argon laser (laser power: 20 W, laser intensity: 1.5-2%) and detected with a spectral PMT (range: 500 nm – 550 nm, gain: 800). The mCherry signal was visualized by exciting with a 561 nm DPSS laser (laser intensity: 1-1.5%) and detected with a spectral HyD (range: 570 nm – 625 nm, gain: 100). Brightfield images were obtained using the transmission PMT detector (gain:275-500, offset: - 2).

### 4.8 Bulk-cell DNA sequencing

For bulk DNA sequencing, bulk NlaIII sequencing was employed following the procedures similar to the single-cell approach [1], nuclei of each cell population (1000 cells, amplified from CIN-seq-collected single-cell clones, 1 Control and 6 TP clones) underwent restriction enzyme digestion, resulting in DNA fragments that were ligated to barcoded adapters. These ligated fragments were then subjected to in vitro transcription and prepared as sequencing libraries, followed by 150 bp paired-end sequencing using the Illumina NextSeq 2000 platform at a mean coverage rate of 70M of mapped reads. The raw fastq files obtained from the sequencing were aligned against the human genome assembly (GRCh38) using the Burrows-Wheeler aligner (BWA) [29]. Subsequently, mapped data underwent further analysis, including the removal of reads lacking a NlaIII sequence and PCR-duplicated reads. Copy number analysis of bulk DNA sequencing data was performed using the AneuFinder R package (v.1.26.0) [25]. Default parameters were employed, including the Edivisive method with a fixed bin size of 1 Mb. Additionally, a blacklisting strategy was implemented to exclude artifacts stemming from high-repeat regions and to correct GC content for CNV calling. The genomes were divided into non-overlapping intervals of varying size based on mappability, with an average of 1 Mb. CNV profiles were visualized as clustergrams using Matlab.

### 4.9 Fluorescence in situ hybridization (FISH)

The cells were initially subjected to time-lapse recording using the UFO microscope. Following the recording, the cells underwent fixation with Carnoy’s solution (60% ethanol, 30% chloroform, 10% acetic acid) after a single wash with PBS. Following a 10-min incubation period, the fixative was removed to allow the cells to dry. Subsequently, the cells were treated with 1 mL of 2X saline-sodium citrate (SSC) medium, followed by sequential washes with 70%, 80%, and 100% ethanol. Next, 3 μL of Chromosome 6 or 16 enumeration probes (CytoCell), pre-mixed with 7 μL hybridization solution, were added to the cells. The cells were covered with a coverslip, sealed with nail polish, and subjected to incubation at 37°C for 5 min followed by 2 min at 75°C for denaturation. After denaturation, the cells, along with the probe, were hybridized at 37°C for 1 hr in a humidified container. Subsequently, the cells were washed with 0.25X SSC and 2X SSC containing 0.05% Tween-20. Finally, the cells were stained with DAPI antifade.

### 4.10 Validation assays using small compounds

Two days before conducting experiments, approximately 100,000 cells (MCF10A or MCF7) were seeded on a fibronectin-coated 6-well glass-bottom plate (Cellvis) in culture medium supplemented with 1X SPY650-DNA nuclear stain. After allowing the cells to attach for one day, the medium was supplemented with 9 μM RO-3306 (Sigma Aldrich, SML0569) to synchronize cells’ cell cycle phase before the G2/M stage. For the validation of RhoGTPase Cycle signaling, 0.05 μg/mL of Rho/Rac/Cdc42 Activator I (Cytoskeleton, CN04) was incubated with the cells for 4 hr. To validate the PTEN signaling, 1.4 μM bpV(HOpic) (Sigma-Aldrich, SML0884) was incubated with the cells 12for 17 hr. For the validation of the involvement of the BCL2L1 gene in promoting cell survival, A-1331852 (Tocris Bioscience, 7661), a BCL2L1 inhibitor, was administered at a concentration of 0.5 nM to the cells and incubated for 4 hr. Following the incubation period, the cells were washed three times with pre-warmed PBS and then incubated with new culture medium without RO-3306. Subsequently, the cells underwent live-cell, multi-region, time-lapse recording using the UFO microscope. To investigate the functional roles of degranulation-associated lysosomal enzymes and anti-apoptotic pathways in tripolar (TP) division cells, MCF10A cells were treated with 2 μM Pepstatin (CTSD inhibitor; MedChemExpress, HY-P0018; 4 μl of a 1 mM stock in 2 ml of MCF10A imaging culture medium) for 24 hr. In additional experiments validating the effect of the anti-apoptotic protein BCL2L1 on CTSD-associated lysosomal death pathways, cells were co-treated with 2 μM Pepstatin for 24 hr and 0.5 nM A-1331852 (BCL2L1 inhibitor) for 4 hr. Real-time mitosis detection was then performed following each treatment to assess the survival rates of daughter cells produced by multipolar divisions, using fresh culture medium without RO-3306. To assess cytokinesis failure, cells undergoing mitosis were monitored 12 hr post-mitosis, and instances where multiple nuclei merged within single cells were identified as indicative of failed cytokinesis. For evaluating cell survival rates post-mitosis, we tracked the viability of cells 12 hr after the completion of mitosis.

### 4.11 Immunoblot

For total protein extraction, cells were lysed in RIPA buffer (ab156034, Abcam) supplemented with Halt™ Protease and Phosphatase Inhibitor Cocktail (78440, Thermo Fisher Scientific). Equal amounts of protein were denatured by boiling with 4× Laemmli loading buffer at 100 °C for 5 min and then resolved by SDS-PAGE using 4–12% Bis-Tris gels (NP0322BOX, Invitrogen). Following electrophoresis, proteins were transferred onto polyvinylidene difluoride (PVDF) membranes (IPVH00010, Millipore). To block nonspecific binding, membranes were incubated in 5% skim milk prepared in TBS containing 0.1% Tween-20 (TBS-T) for 2 h at room temperature. The membranes were subsequently incubated overnight at 4 °C with primary antibodies against Akt (pan) (#2920S, 1:1000, Cell Signaling Technology), phospho-Akt (#4060S, 1:1000, Cell Signaling Technology), GAPDH (#2118T, 1:1000, Cell Signaling Technology), and Rho (#8789, 1:1000, Cell Signaling Technology). After washing, membranes were incubated with the appropriate HRP-conjugated secondary antibodies for 2 h at room temperature. Protein bands were visualized using the ECL™ Prime Western Blotting Detection System (GERPN2232, Sigma) and imaged with the Amersham imaging system.

The Active Rho Detection Kit (#8820, Cell Signaling Technology) was employed to assess the active form of RhoA, following the manufacturer’s protocol in conjunction with immunoblotting. Briefly, cells were collected, washed with ice-cold PBS, and lysed in lysis buffer. Cell lysates were centrifuged at 16,000 × g for 15 min to remove debris. Agarose beads pre-equilibrated with glutathione resin were added to spin cups with collection tubes and incubated with GST-Rhotekin-RBD. The clarified cell lysates were immediately transferred to the spin cups and incubated for 1 h at 4 °C with gentle rotation to allow active RhoA binding to the resin. The spin cups were then centrifuged at 6,000 × g for 30 s to separate the beads from the supernatant. After washing, the samples were mixed with reducing sample buffer containing dithiothreitol (DTT) and incubated for 2 min at room temperature. The tubes were centrifuged again at 6,000 × g for 2 min. Finally, the eluted proteins were heated at 100 °C for 5 min and subjected to immunoblotting.

### 4.12 Lentivirus production and transduction

Lentiviral particles were generated using HEK293 cells. One day before transfection, HEK293 cells were seeded in 10 cm culture dishes at approximately 50% confluency. Transfection was performed when the cells reached 70–80% confluency, typically 24 hours after seeding. For each 10 cm dish, a plasmid mixture was prepared containing 8.5 μg of the vector of interest (#116780, pHAGE-PTEN; #98323, BCL2L1_pLX307, Addgene), 4.5 μg of psPAX2 (second-generation lentiviral packaging plasmid, #12260, Addgene), and 4.5 μg of pMD2.G (VSV-G envelope-expressing plasmid, #14888, Addgene). Transfection was performed using Lipofectamine 3000 reagent (L3000150, Thermo Fisher Scientific) following the manufacturer’s protocol. Briefly, 53 μL of Lipofectamine 3000 reagent and 35 μL of P3000 Enhancer were mixed with the plasmid DNA in 2.5 mL of Opti-MEM medium (Gibco).

Viral supernatants were harvested 24 and 48 hours post-transfection and filtered through a 0.45 μm PVDF membrane (Sigma-Aldrich) to remove cell debris. The viral particles were subsequently concentrated by ultracentrifugation using a Beckman Coulter ultracentrifuge. The concentrated viral stock was aliquoted and stored at −80 °C until further use.

For transduction, MCF-10A cells were seeded in 6-well plates at approximately 60% confluency. After 24 hours, the culture medium was replaced with fresh medium containing 10 μg/mL Polybrene (TR-1003, Millipore) to enhance infection efficiency. Following another 24-hour incubation, the virus-containing medium was replaced with fresh complete culture medium. Puromycin selection was initiated 48 hours post-transduction using 1.5 μg/mL Puromycin (A1113803, Thermo Fisher Scientific).

### 4.13 Proliferation assay

Proliferation assays were performed using the Cell Counting Kit-8 (96992, Sigma-Aldrich). 100 μl cell suspensions with medium was added to a 96-well plate(2 × 103 cell per well) and incubated for 12 hour at 37°C. Then, 10 μl CCK-8 reagent was added to each well, cells were incubated 3 hours at 37°C and optical density at 450 nm was measured. Proliferation assays were performed after 24, 48 and 72 hours. The growth rate was expressed as a value relative to hour 0.

### 4.14 Statistical analysis

The number of cells analyzed and preprocessing normalization of data are indicated in the corresponding figure legends. Data were analyzed using R software (version 4.3.3) or Microsoft Excel. Results are expressed as mean ± standard deviation (SD) when referring to the variability within the entire population (Excel function: STDEV.P). When applicable, the standard error of the mean (SEM) was calculated to indicate how much the sample means are expected to vary from the true population mean. In Excel, SEM was obtained as =STDEV.S(range)/SQRT(COUNT(range)). The specific measure used (SD or SEM) is indicated in the corresponding figure legends. For statistical analysis, Student’s t-tests (for two comparisons) or Wilcoxon test were used to compare groups in R, as appropriate. The overlaid quantiles (25th, 50th, 75th percentiles) represent the data spread. Individual data points are shown as jittered points. Bonferroni correction is applied to adjust the p-values for multiple comparisons to control the false positive rate.

The differences in single-cell RNA omics data were calculated using limma that apply an empirical Bayes method to moderate the standard errors of the estimated log-fold changes. The Ingenuity Pathway Analysis (IPA) primarily uses Fisher’s exact test to evaluate the statistical significance of pathway enrichment.

For statistical analyses, the ggpubr (v0.6.0) and dplyr (v1.1.4) packages were employed. A *p*-value less than 0.05 was considered statistically significant. Data values: ns (not significant); p < 0.05 *; p < 0.01 **; p < 0.001 ***; p < 0.0001 ****.

### 4.15 Data Visualization

Dot plots, violin plots, and histograms were generated using the ggplot2 package (v4.0.0). Volcano plots were created with the EnhancedVolcano package (v1.20.0). Heatmaps were constructed using the ComplexHeatmap package (v2.18.0)), and Seurat package (v4.4.0). Bar plots were generated in Microsoft Excel. Additional visualization and data handling packages included RColorBrewer (v1.1-3), patchwork (v1.2.0), and tidyverse/readxl (v1.4.3). Pathway analyses were conducted using Ingenuity Pathway Analysis (IPA, Q2 2025), and schematic illustrations were created with PowerPoint (Microsoft 365. Windows 10).

## Data Availability Statement

The main data supporting the results in this study are available within the paper and its xxxx Data generated in this study, including source data and the data used to make the figures, will be available from Open Science Framework at xxx. The sequencing data will be available at the NCBI BioProject database with the BioProject ID xxx. The sequencing data analysis used in the study will be available on GitHub before publication.

## Acknowledgements

M.-P.C. acknowledges the support from the ERC Consolidator Grant, Oncode Institute, NWO (the Netherlands Organization for Scientific Research) Vidi and Aspasia Grants, Stichting Ammodo Grant, and Erasmus MC-TU Delft-EUR Flagship Grant. We thank the Josephine Nefkens Cancer Program for their infrastructural support. P.-R.S. acknowledges the PhD fellowship support provided by the Ministry of Science and Technology in Taiwan. We thank Reza Ghadiri Rad for his assistance for the validation assays; R. Agami and Maayke Kuijten for the kind gift of MCF10A and MCF7 cell lines, respectively; S. Lens, G. Kops and R. Medema, for discussions about the project.

## Author contributions

P.-R.S. and T.-C.C. designed and conducted experiments, performed image analysis, and contributed to data interpretation. P.-R.S. and M.T.L.-C. developed scripts and conducted analysis of the single-cell DNA and RNA sequencing data. S.v.W. performed the CRISPR imaging assay and analysis. C.L. scripted part of the code for and contributed to the analysis of single-cell DNA sequencing data. C.B. and T.-Y.C. assisted in cell culture preparations and validation assays. L. Y. assisted in image analysis and cell tracking. J.S. provided technical support with the UFO microscope. M.-P.C., P.-R.S., and T.-C.C. wrote the manuscript with input from all authors. M. P.C. initiated, contributed to, and supervised all aspects of the project.

## Supporting Information

### Characteristics of MCF10A TP cells

We observed a high survival rate among TP cells (99.4%, defined as surviving for > 24 hr or until the next mitosis). Among the surviving, mitotic TP cells, 63.2% continued with tripolar mitosis, while 36.8% reverted to bipolar mitosis (Figure S4a-ii and S4b; Figure S5). These results suggest the phenotypic plasticity of TP cells, as evidenced by their successful survival in the population and their ability to transition between mitosis types over generations.

Regarding the mother cells of TP cells, the majority of TP cells originated from mother cells undergoing bipolar mitosis (74.1%) (Figure S4a-ii), while others originated from mother cells undergoing tripolar/multipolar mitosis (25.9%) (Figure S4a-ii). Notably, 50.3% of TP cells originated from binucleate cells, where two nuclei merged or fused within a single cell (Figure S4c, S4d). This phenomenon may be attributed to cytokinesis failure or segregation errors over time.

## Supporting Figures

**Figure S1.**
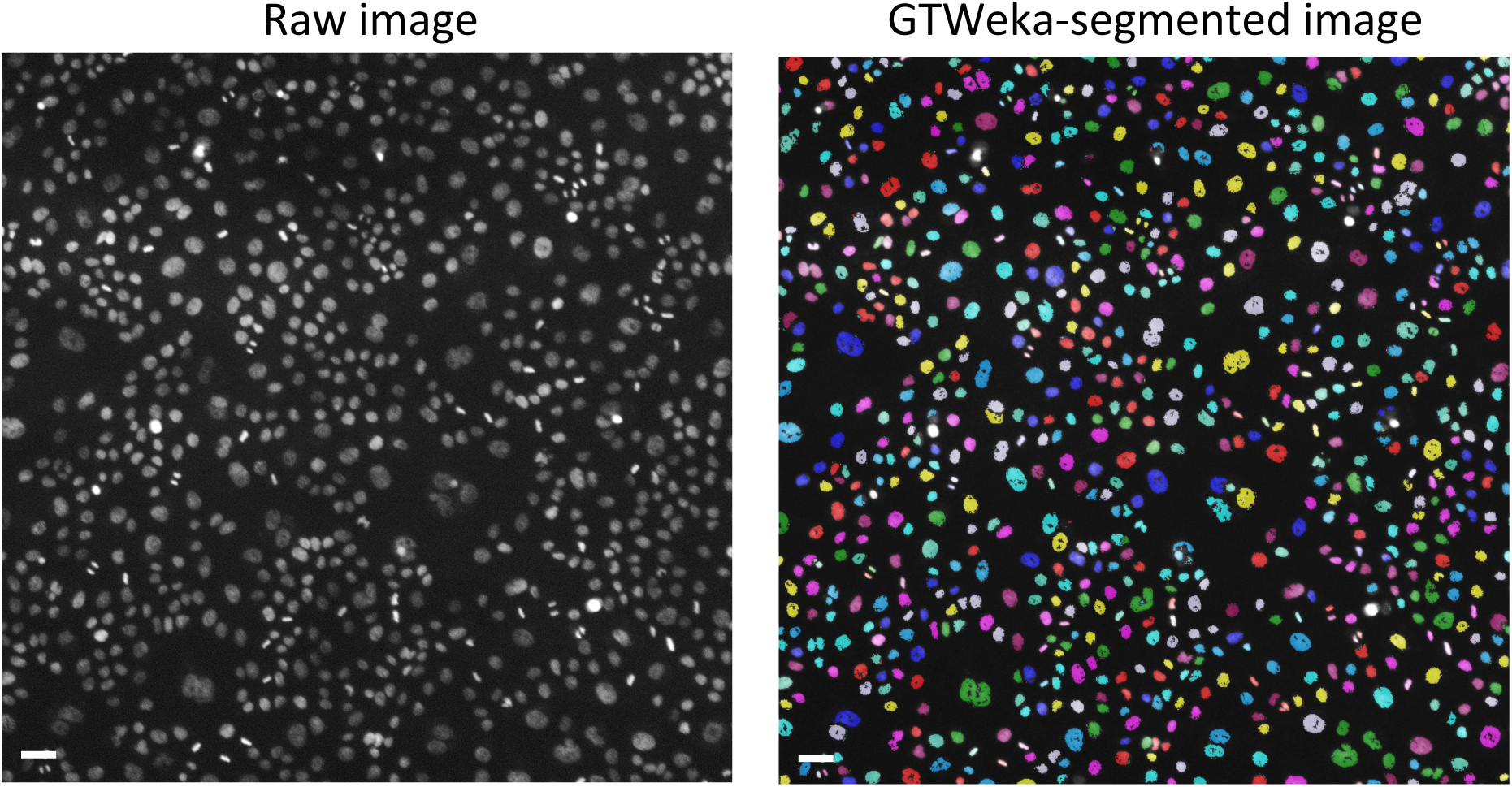
GTWeka cell segmentation. Real-time cell segmentation within CIN-seq is achieved through the ground-truth assisted trainable Weka segmentation (GTWeka) method, integrated within the CIN-seq pipeline. Scale bars denote 50 μm.

**Figure S2.**
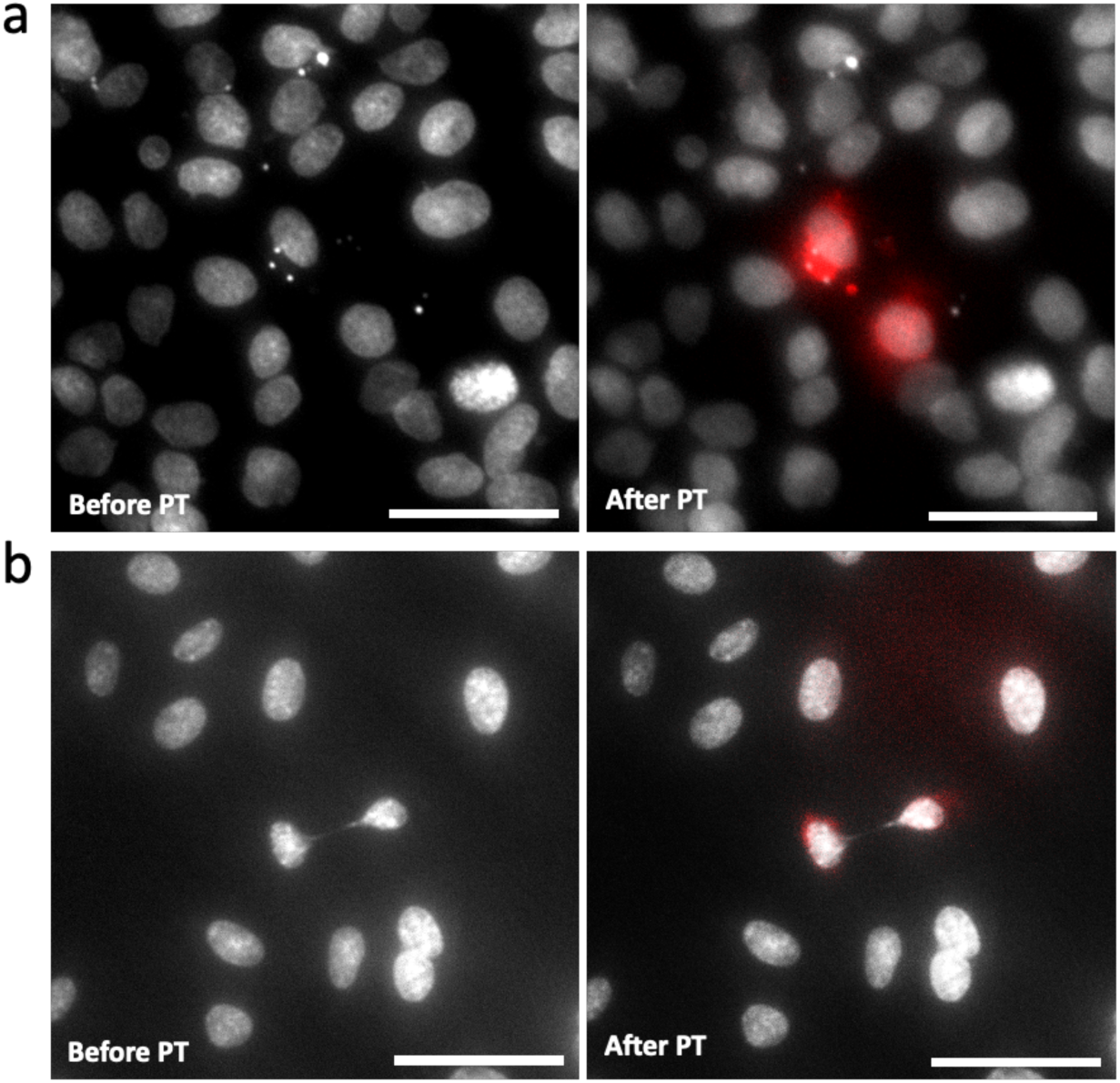
Additional CIN phenotypes. a) A representative image shows two instances of micronucleated (MN) cells, products of lagging chromosomes, from patient-derived esophageal adenocarcinoma (EAC) cells stained with SPY650-DNA nuclear dyes. Left panel: the image before phototagging (PT). Right panel: the image after PT (red). b) A representative image shows one instance of a chromatin bridge cell from patient-derived EAC cells stained with SPY650-DNA nuclear dyes. Left panel: the image before PT. Right panel: the image after PT (red). The scale bars denote 100 µm.

**Figure S3.**
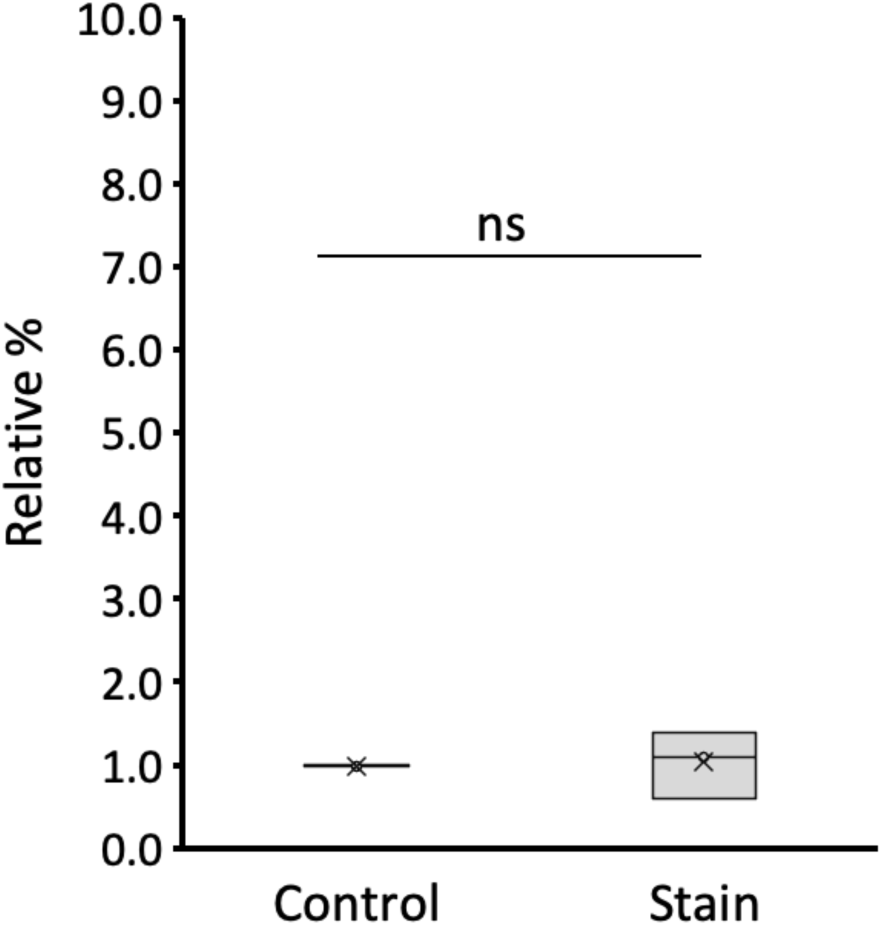
Bar plot showing the ratio of cells undergoing tripolar mitosis versus bipolar mitosis under two conditions: Control (no dyes added) and Stain (both SPY650 DNA dye and photoactivatable dye added). ∼2,700-2,900 mitotic events were analyzed in both conditions (N = 3). The p-value was obtained using Student’s t-test. ns (not significant).

**Figure S4.**
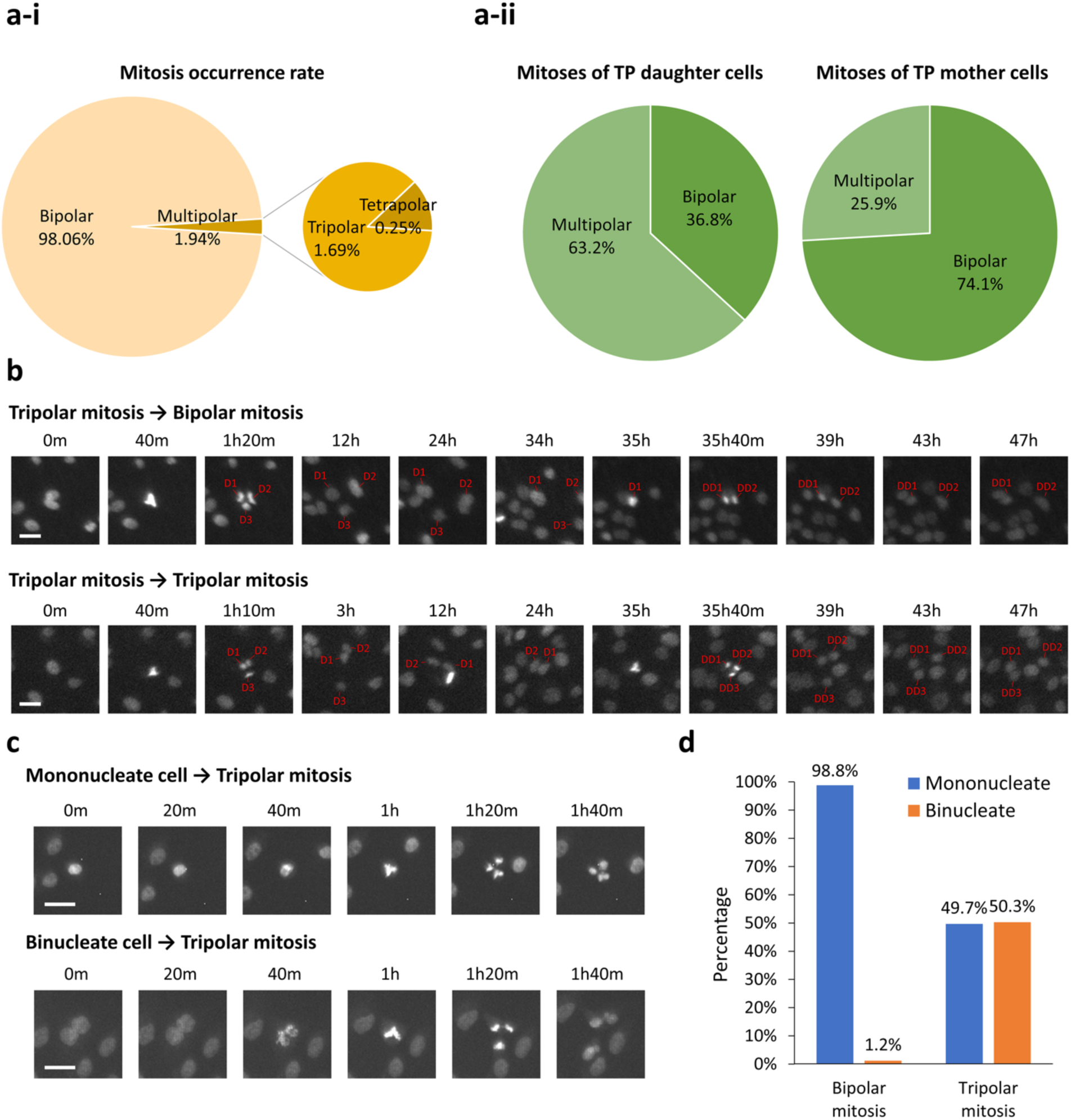
Quantification of multipolar mitoses in MCF10A cells. a-i) Statistics of the rates of bipolar and multipolar mitoses in MCF10A cells. a-ii) Statistics of mitosis phenotypes for TP daughter cells and mother cells. b) Representative time-lapse images of tripolar daughter cell(s) followed by either bipolar mitosis (upper panels) or tripolar mitosis (lower panels). c) Representative time-lapse images of tripolar daughter cells dividing from mononucleate cells (upper panels) or binucleate cells (lower panels). d) Statistics of the rates of bipolar or tripolar mitoses originating from mononucleate cells or binucleate cells. All the scale bars denote 30 μm.

**Figure S5.**
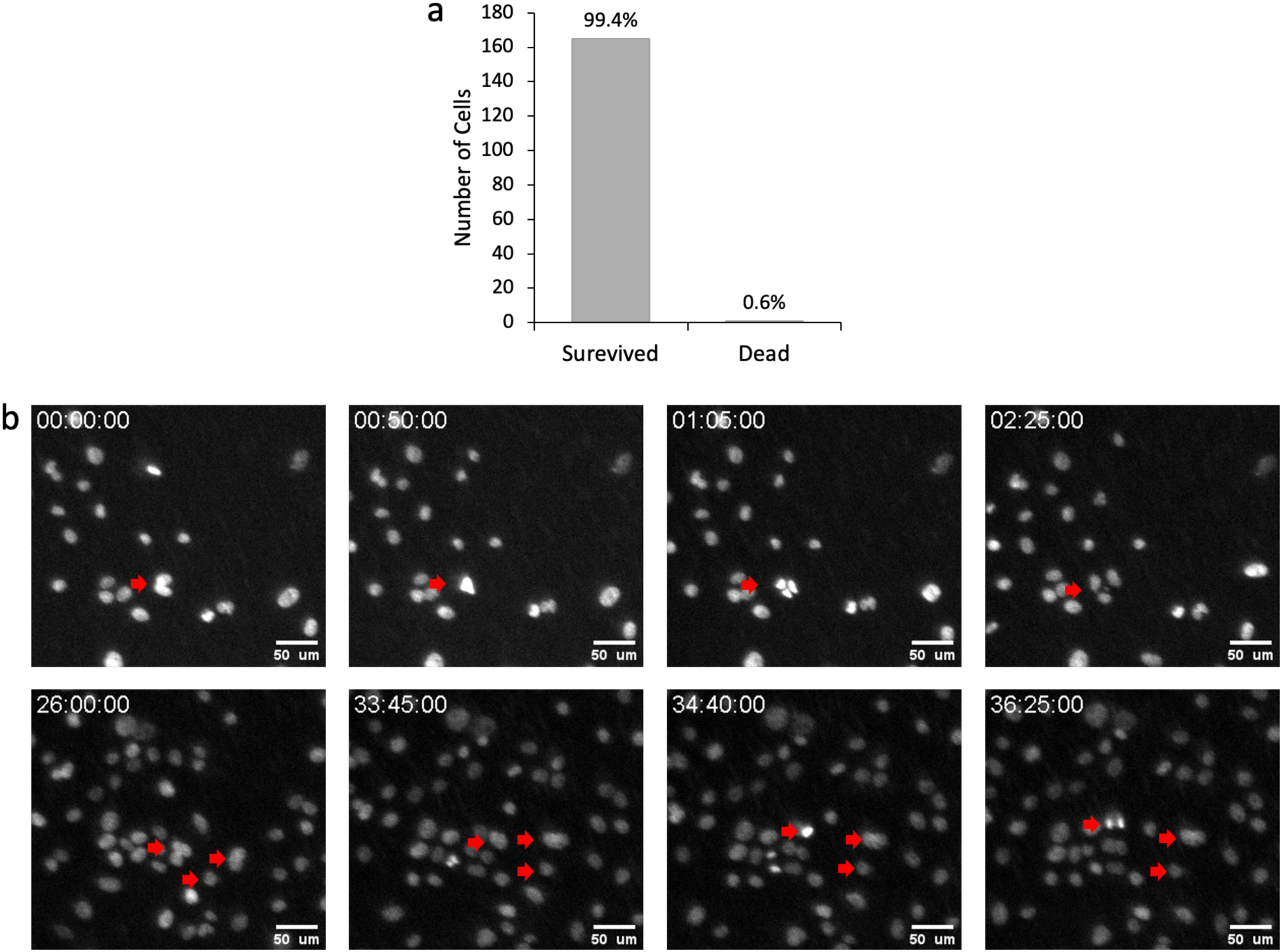
Survival rate of cells after tripolar mitosis (TP cells). a) A total of 166 TP cells were monitored for 24 hours or until the next mitosis; of these, 165 cells survived and 1 cell died. b) Time-course images showing a representative case of tripolar mitosis and the cell fate of the three daughter cells. One of them subsequently underwent bipolar mitosis at 36.5 hours. The scale bars denote 50 µm.

**Figure S6.**
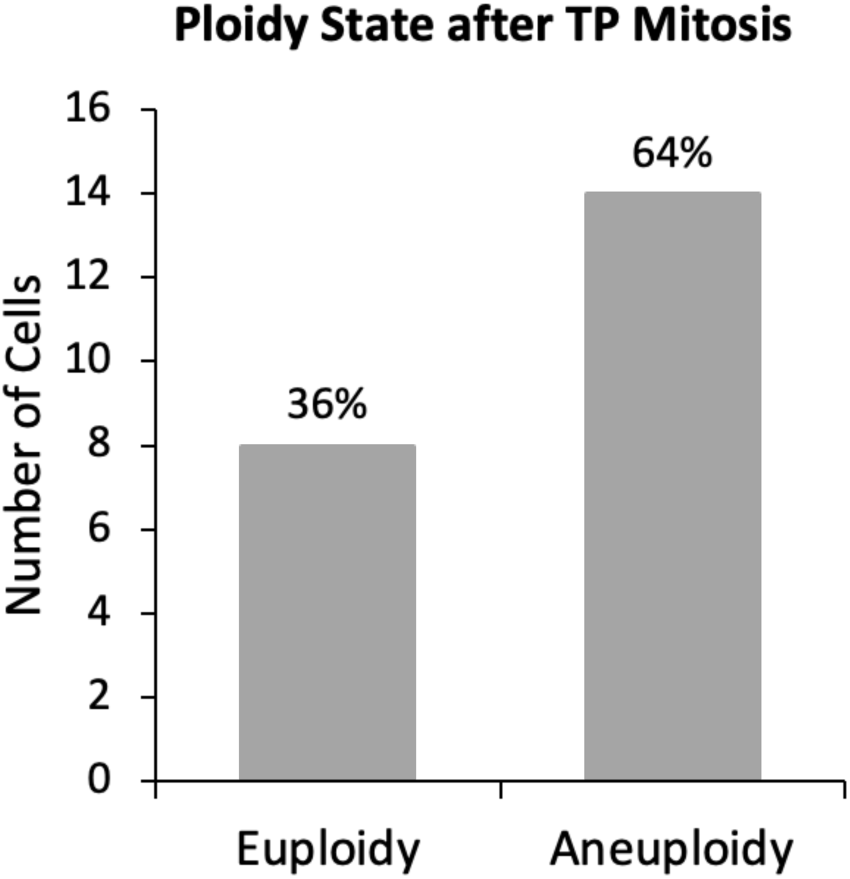
Bar plot showing the number of euploidy or aneuploidy cells after tripolar mitosis.

**Figure S7.**
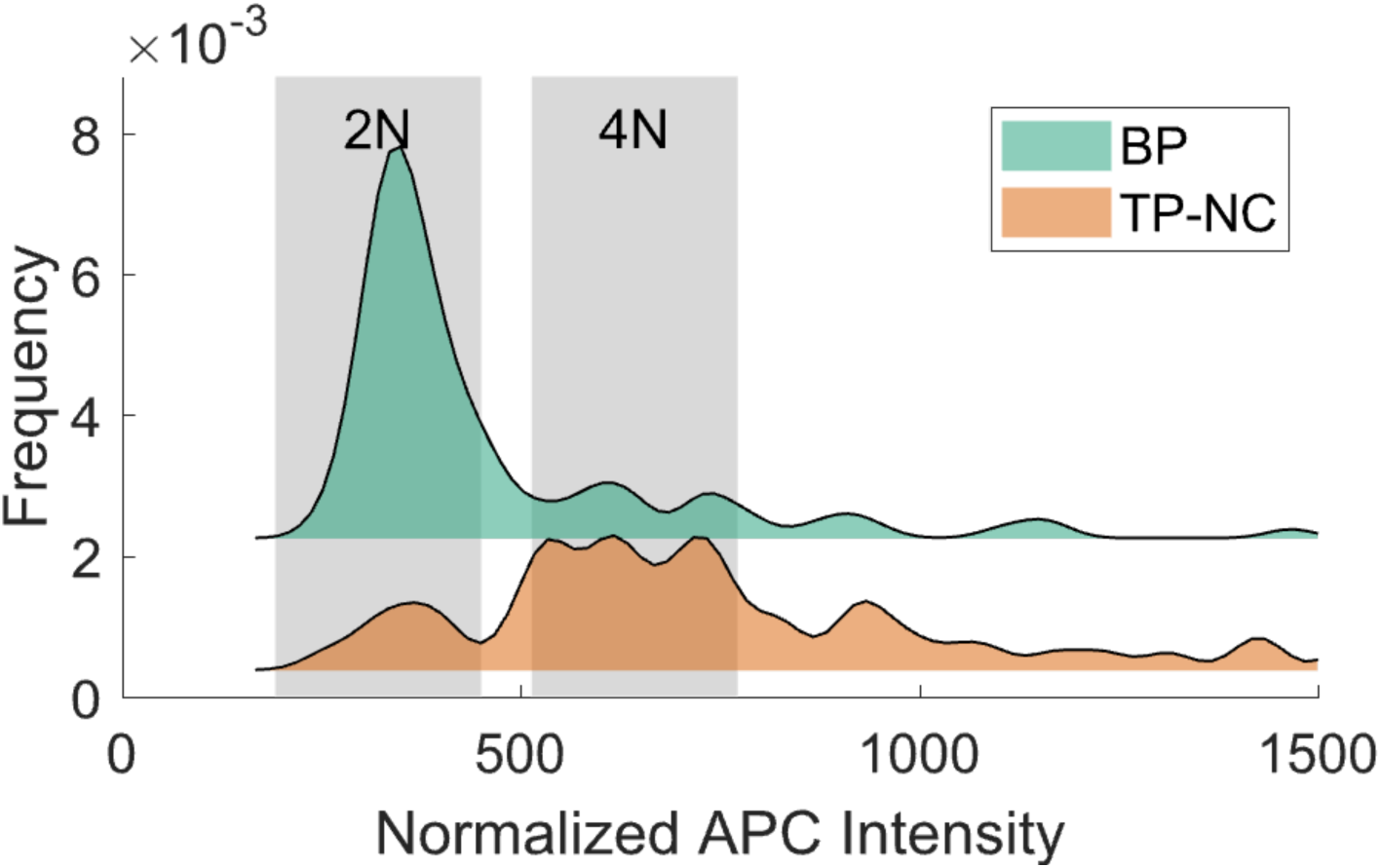
Flow cytometry plots show the DNA content (indicated by APC intensity) of BP and TP-NC cells.

**Figure S8.**
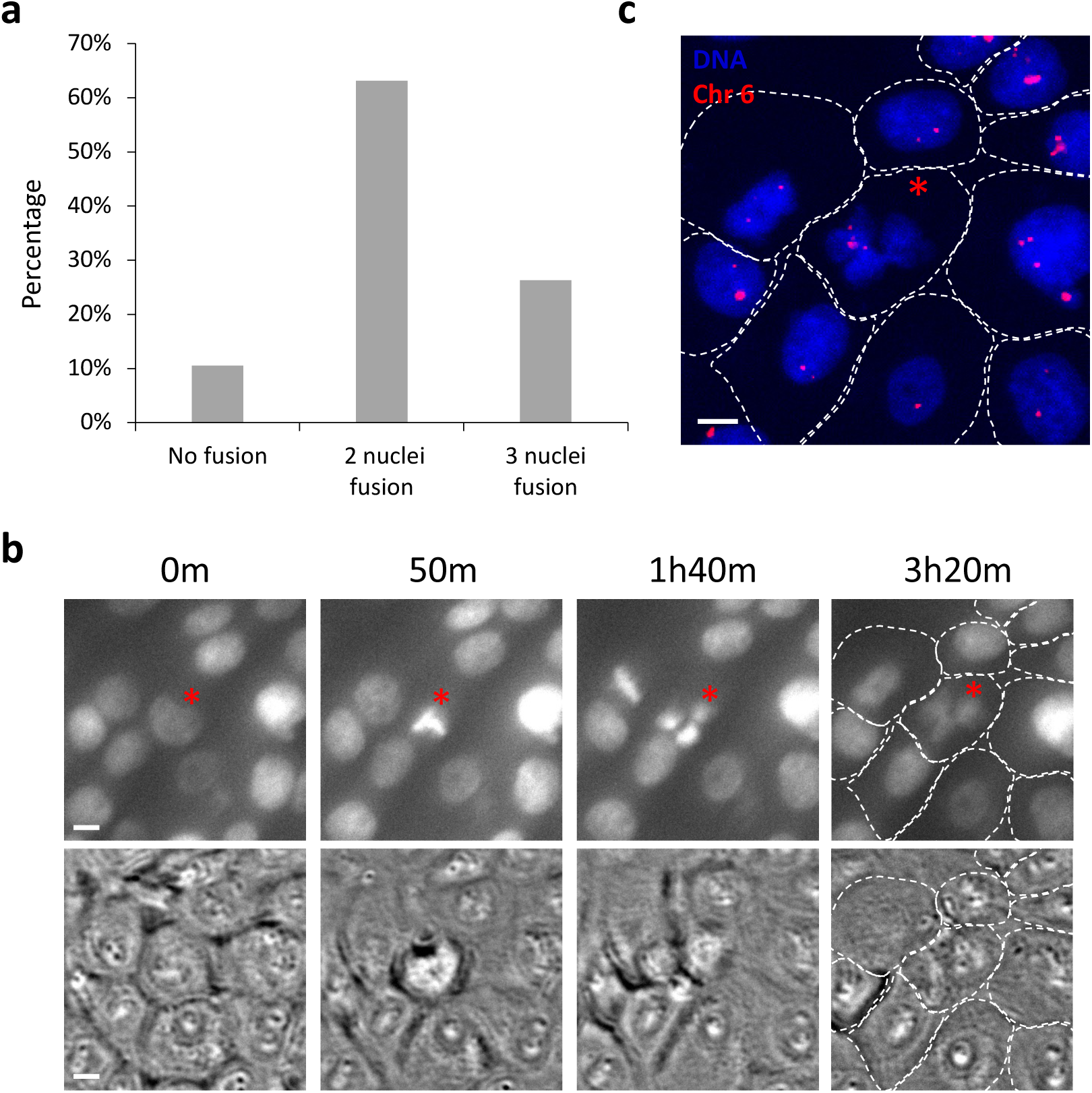
Validation of polyploidy in TP-NC cells. a) Plot of nuclei fusion status, acquired from live-cell time-lapse imaging, within a single cell after tripolar mitosis. ∼65% of cases displayed 2-nuclei fusion, with 2 nuclei aggregating within a single cell, while ∼30% of cases exhibited 3-nuclei fusion, with 3 nuclei merging within a single cell, after tripolar mitosis. b) Representative time-lapse images (bright field (lower panel) and SPY650-DNA nuclear staining (upper panel)) depict cells undergoing 3-nuclei fusion after tripolar mitosis (red asterisk)). Scale bars denote 10 μm. c) FISH assay targeting Chromosome 6 (Chr 6) on fixed cells, corresponding to the last panel of figure e, wherein Chr 6 is visualized in red foci. White dashed lines delineate cell membrane boundaries. The scale bar denotes 10 μm.

**Figure S9.**
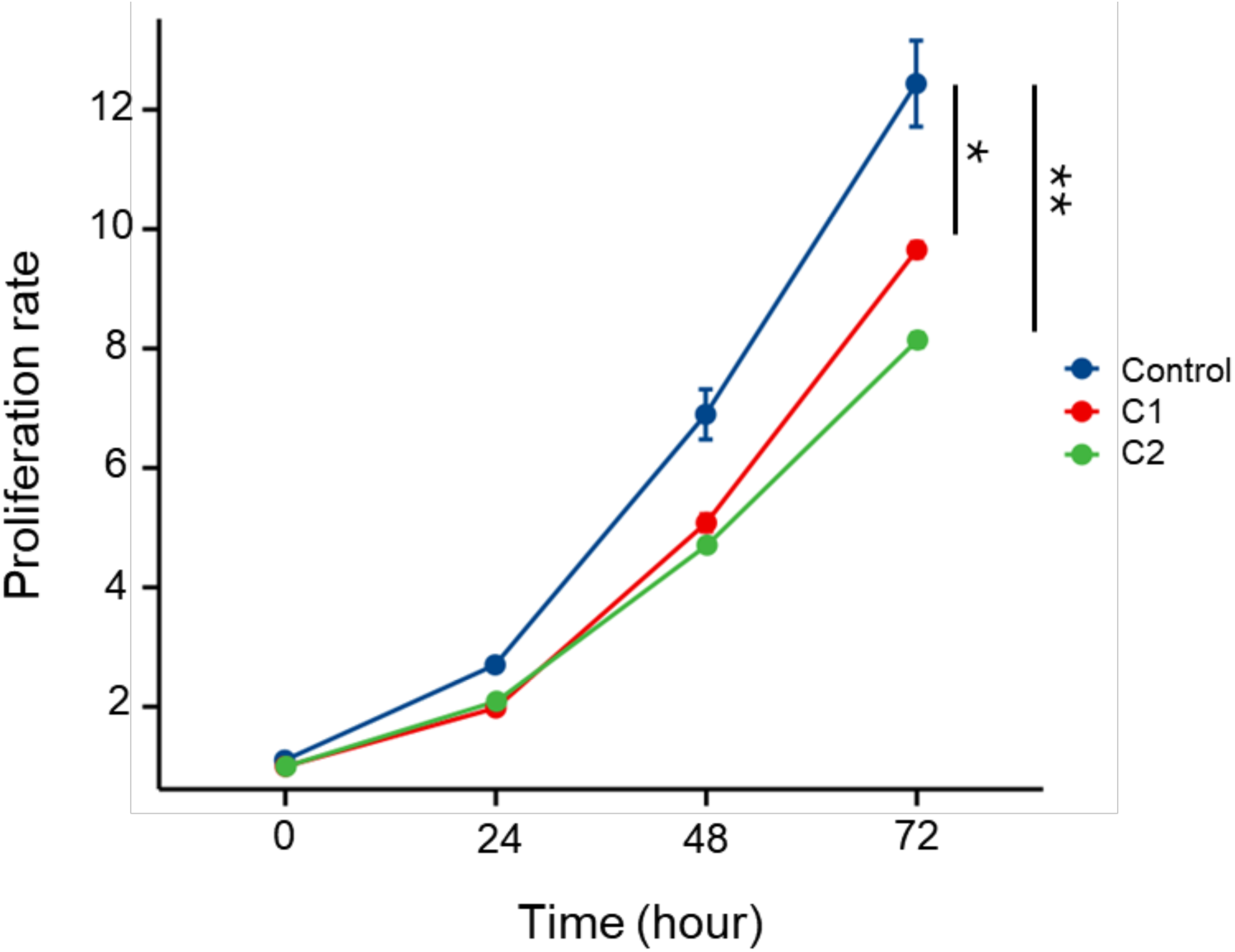
Cell proliferation rate of the bipolar and tripolar cell clones. Cell proliferation rates of the Control bipolar cell clone (blue) and two tripolar cell clones, C1 (red) and C2 (green), were measured at 0, 24, 48, and 72 hours using the CCK-8 assay. Absorbance values were normalized. The proliferation rate increased over time in all groups but was significantly higher in the Control group compared to C1 (p < 0.05 *) and C2 (p < 0.01 **). The p-values were obtained using Two-way repeated-measures ANOVA test.

**Figure S10.**
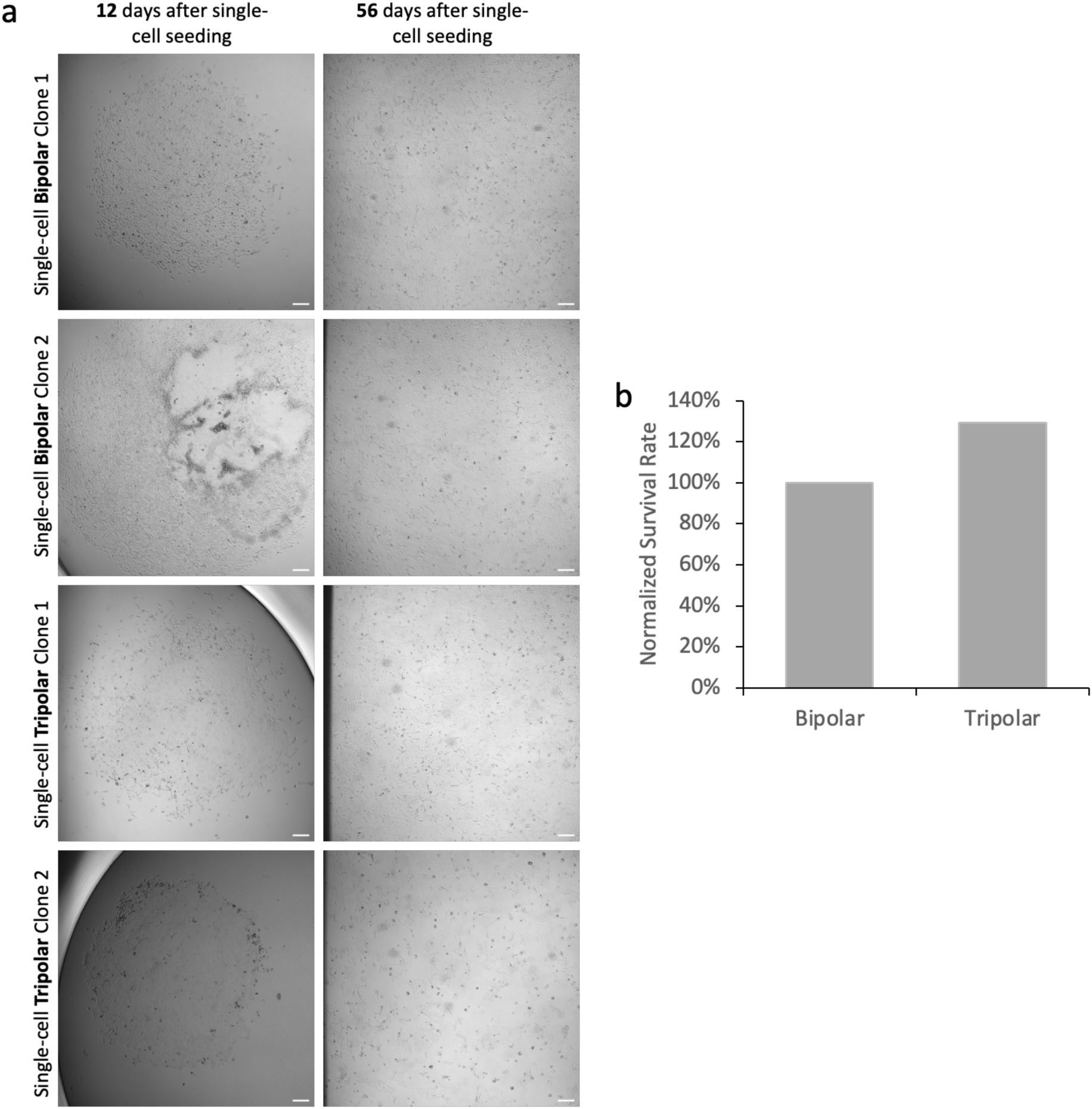
Clonogenic assay. a) Two representative single-cell clones following either bipolar or tripolar mitosis. Images were taken on Days 12 and 56 for these single-cell clones. Scale bars, 200 µm. b) Normalized survival rate (relative to the survival rate of bipolar cell clones). 30 single-cell bipolar clones and 116 single-cell tripolar clones were collected for the assay.

**Figure S11.**
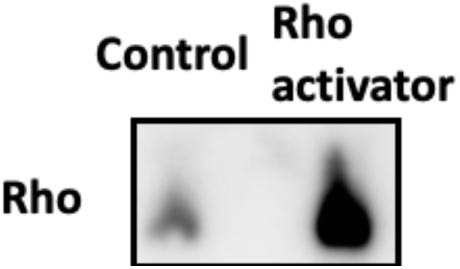
Western blot showing Rho protein expression following Rho perturbation. Western blot analysis showing Rho activation in cells treated with a Rho activator compared with the control group.

**Figure S12.**
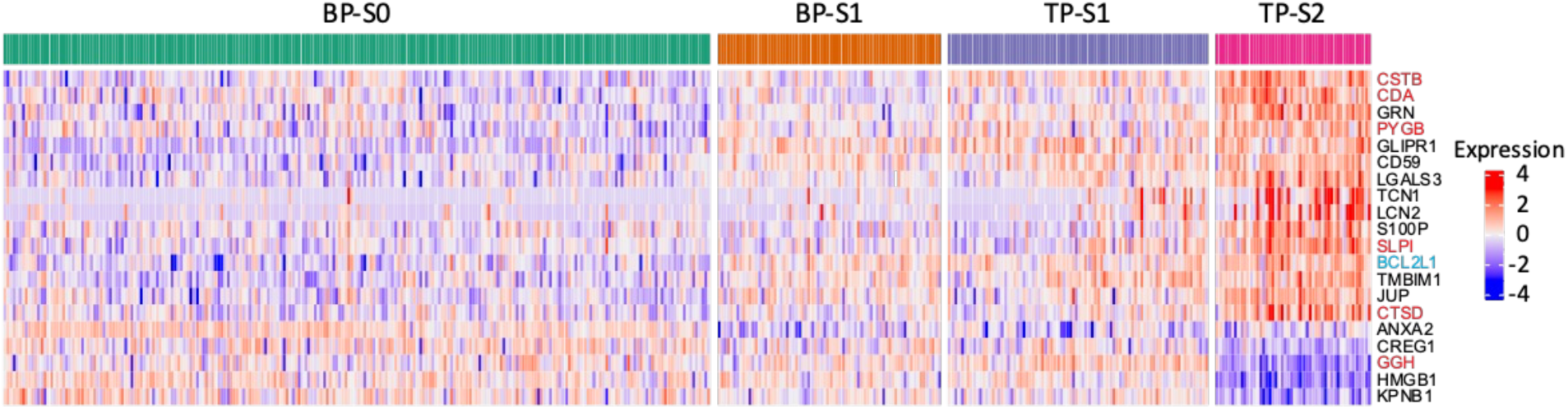
The top 20 differential genes between different cell subclusters (BP-S0, BP-S1, TP-S1, and TP-S2) related to the degranulation-like pathway are shown. Genes related to enzymes or proteases are highlighted in red. The BCL2L1 gene is shown in blue.

**Figure S13.**
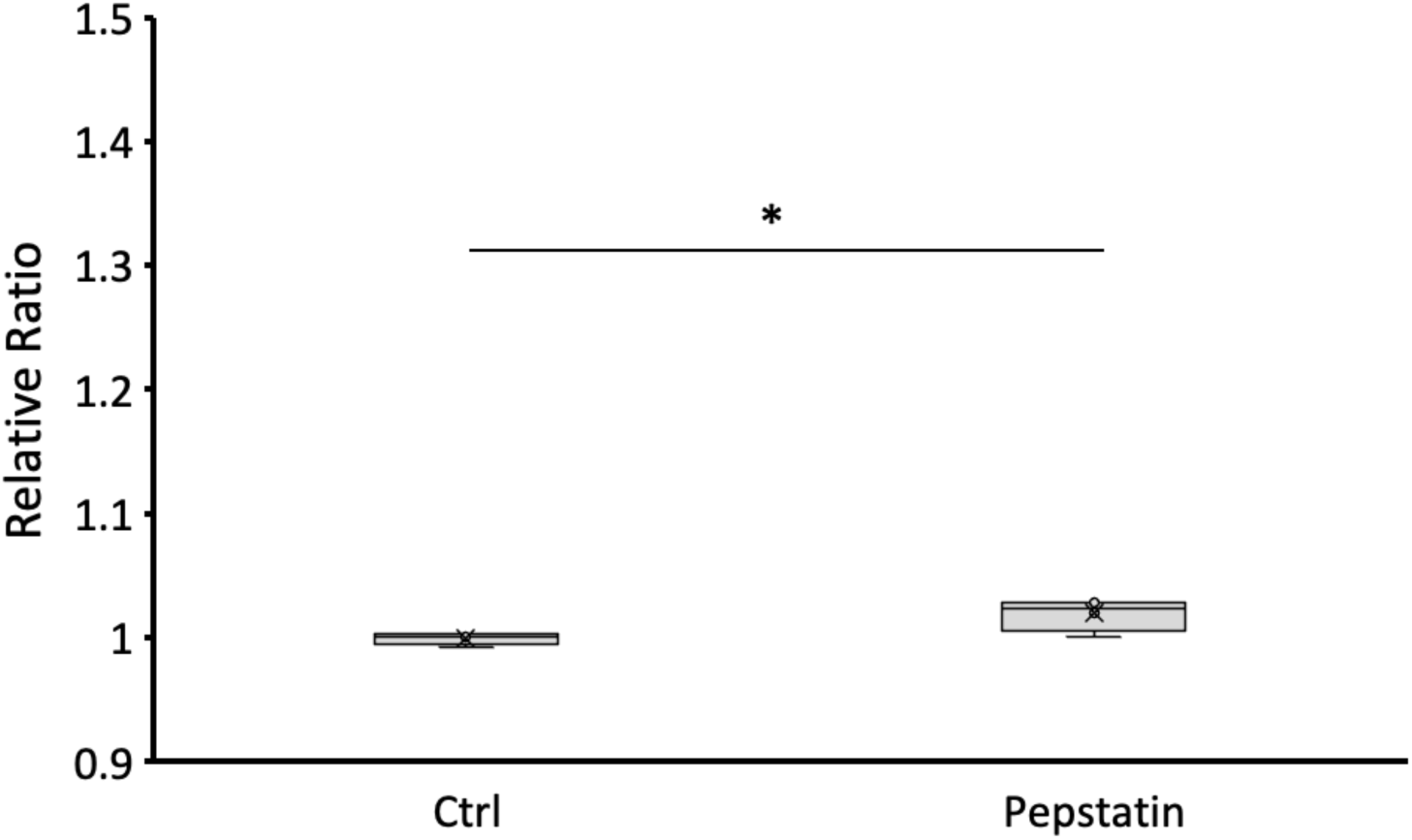
Relative survival rate ratios of TP cells under Control (Ctrl) and Pepstatin-treated, conditions (N = 5). The p-values were calculated using Student’s t-test. p < 0.05 *.

**Figure S14.**
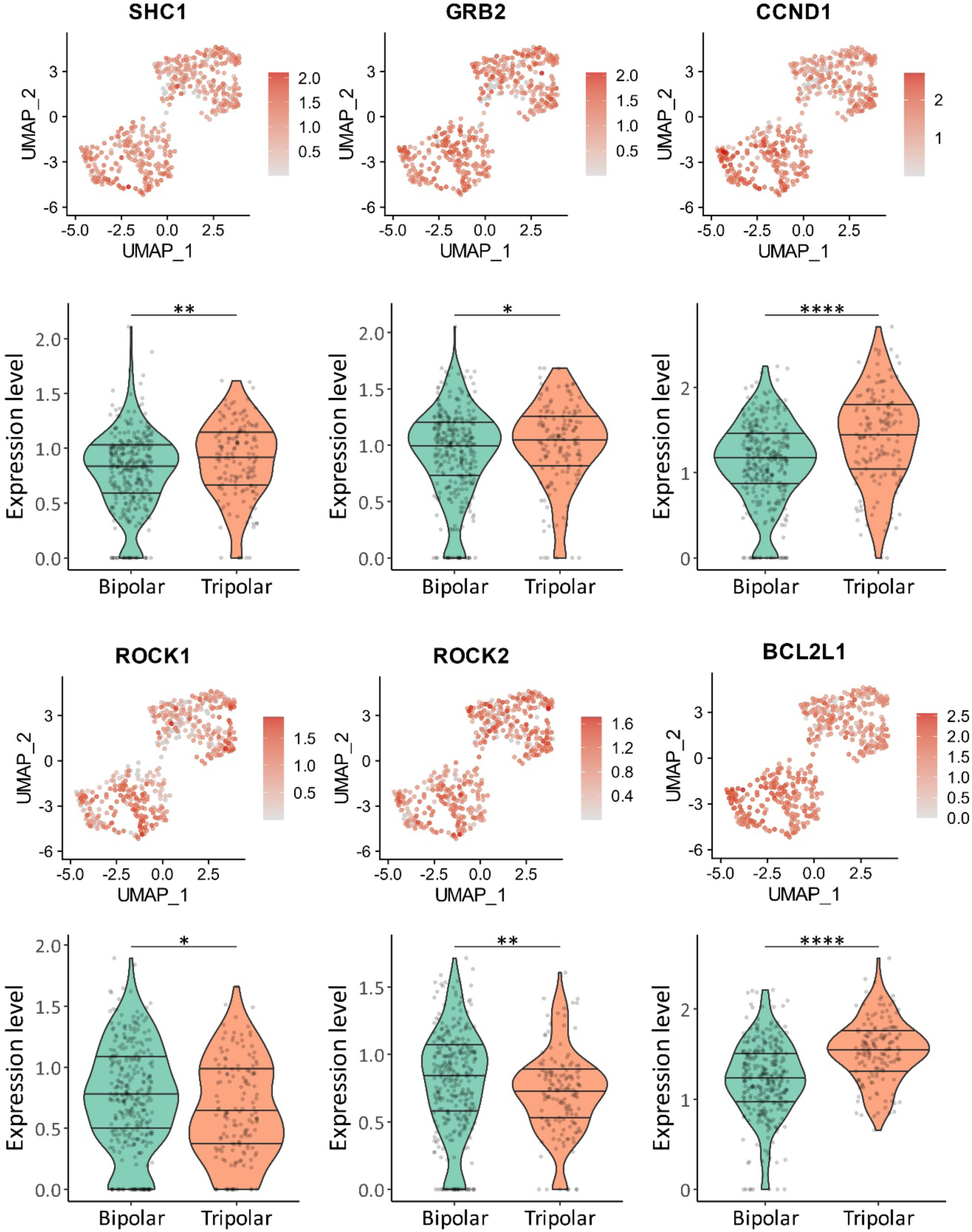
**UMAP plots (upper panels) and violin plots (lower panels) display the module score of genes associated with the PTEN signaling pathway**, specifically SHC1, GRB2, CCND1, ROCK1, ROCK2, and BCL2L1 genes, for bipolar and tripolar cells. The p-value was obtained using Wilcoxon test. Bonferroni correction; p < 0.05 *; p < 0.01 **; p < 0.001 ***; p < 0.0001 ****. The overlaid quantiles (25th, 50th, 75th percentiles) represent the data spread. Individual data points are shown as jittered points. Statistical analysis was carried out using R Software.

**Figure S15.**
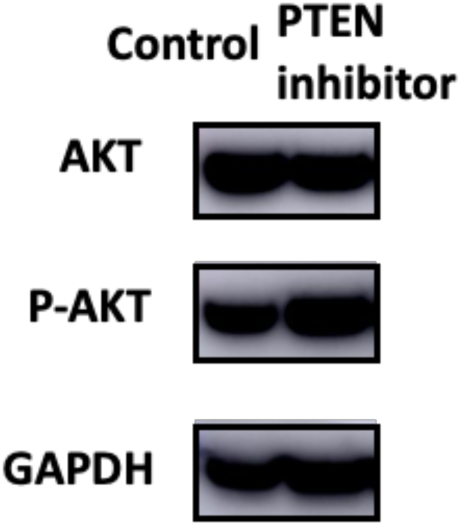
Western blots showing AKT protein expression and phosphorylated AKT following PTEN inhibition. Western blot showing the effects of PTEN inhibition on AKT signaling. Treatment with a PTEN inhibitor increased AKT phosphorylation (p-AKT) without altering total AKT levels. GAPDH was used as a loading control. Please note that PTEN inhibition enhances AKT phosphorylation, while increased PTEN expression suppresses phosphorylated AKT levels.

**Figure S16.**
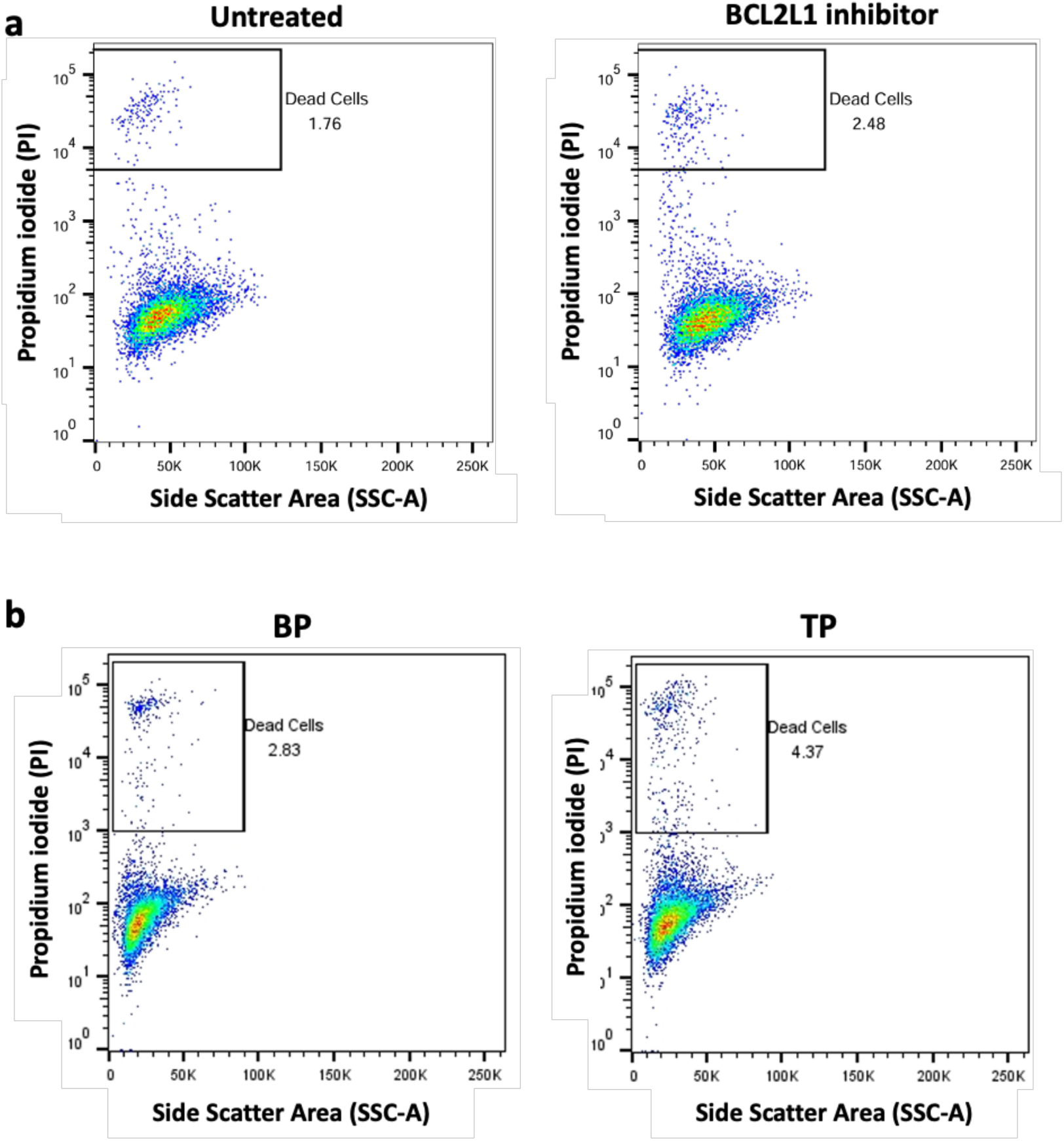
Flow cytometry plots of MCF10A cells following BCL2L1 inhibition. a) left: untreated control; right: cells treated with 0.5 nM BCL2L1 inhibitor (A-1331852; Tocris Bioscience, #7661) for 4 hours. b) Flow cytometry plots of a BP cell clone (left) and a TP cell clone (right) treated with 0.5 nM BCL2L1 inhibitor (A-1331852; Tocris Bioscience, #7661) for four hours. The x-axis represents Side Scatter Area (SSC-A), and the y-axis represents Propidium Iodide (PI) fluorescence, a cell death indicator.

**Figure S17.**
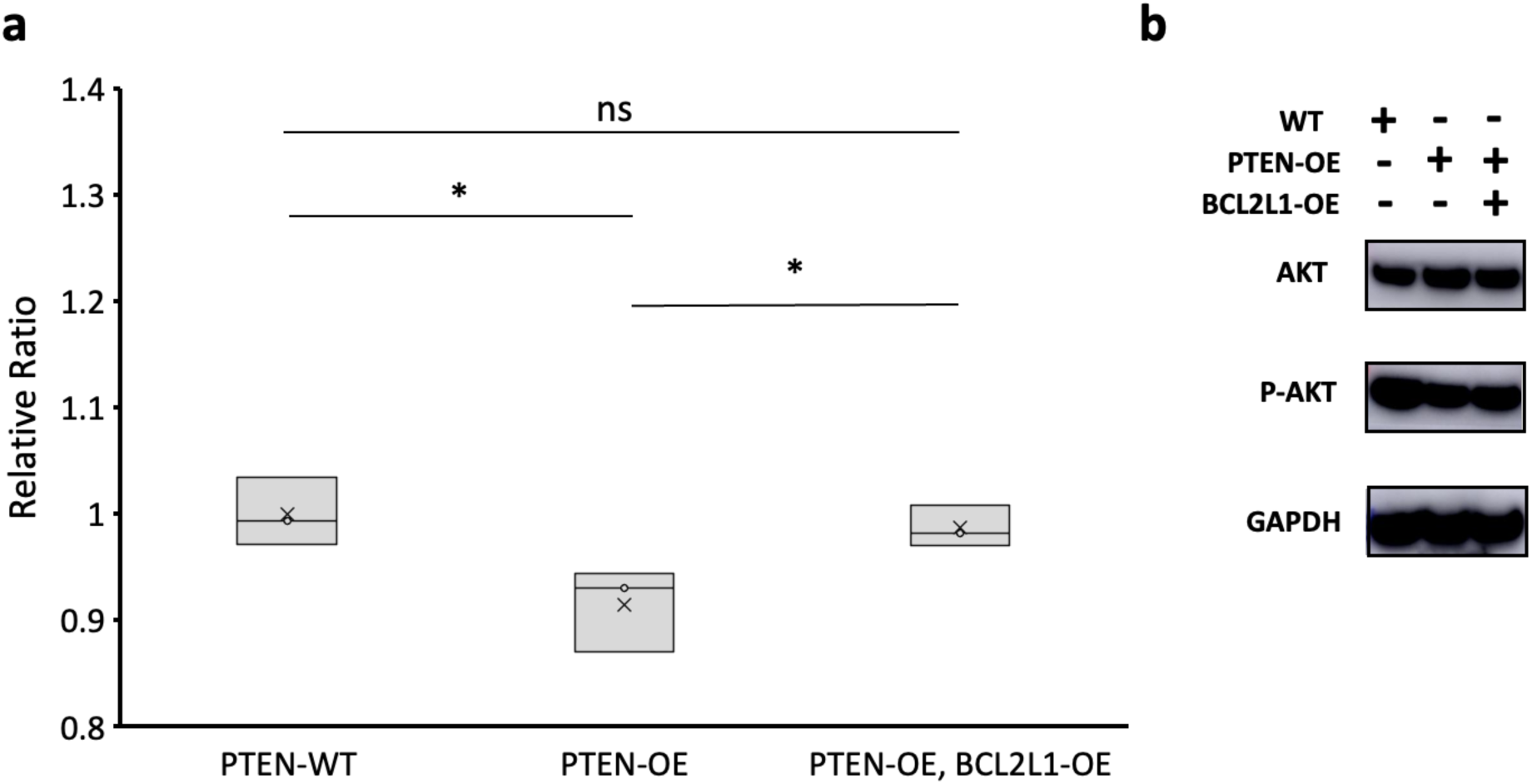
Validation assays with PTEN and BCL2L1 perturbation. a) Box plot shows the normalized survival rate ratios of TP cells in the Control, PTEN-overexpression (OE), and PTEN-OE/BCL2L1-OE groups (N = 3). The p-values were calculated using Student’s t-test. ns (not significant); p < 0.05 *. b) Western blot analysis shows total and phosphorylated AKT (p-AKT; a functional indicator of PTEN inhibition) levels in cells overexpressing PTEN (PTEN-OE) or co-overexpressing PTEN and BCL2L1 (BCL2L1-OE). Overexpression of PTEN decreased AKT phosphorylation, whereas co-overexpression of BCL2L1 partially restored p-AKT levels. GAPDH was used as a loading control. Please note that PTEN inhibition enhances AKT phosphorylation, while increased PTEN expression suppresses phosphorylated AKT levels.

**Figure S18.**
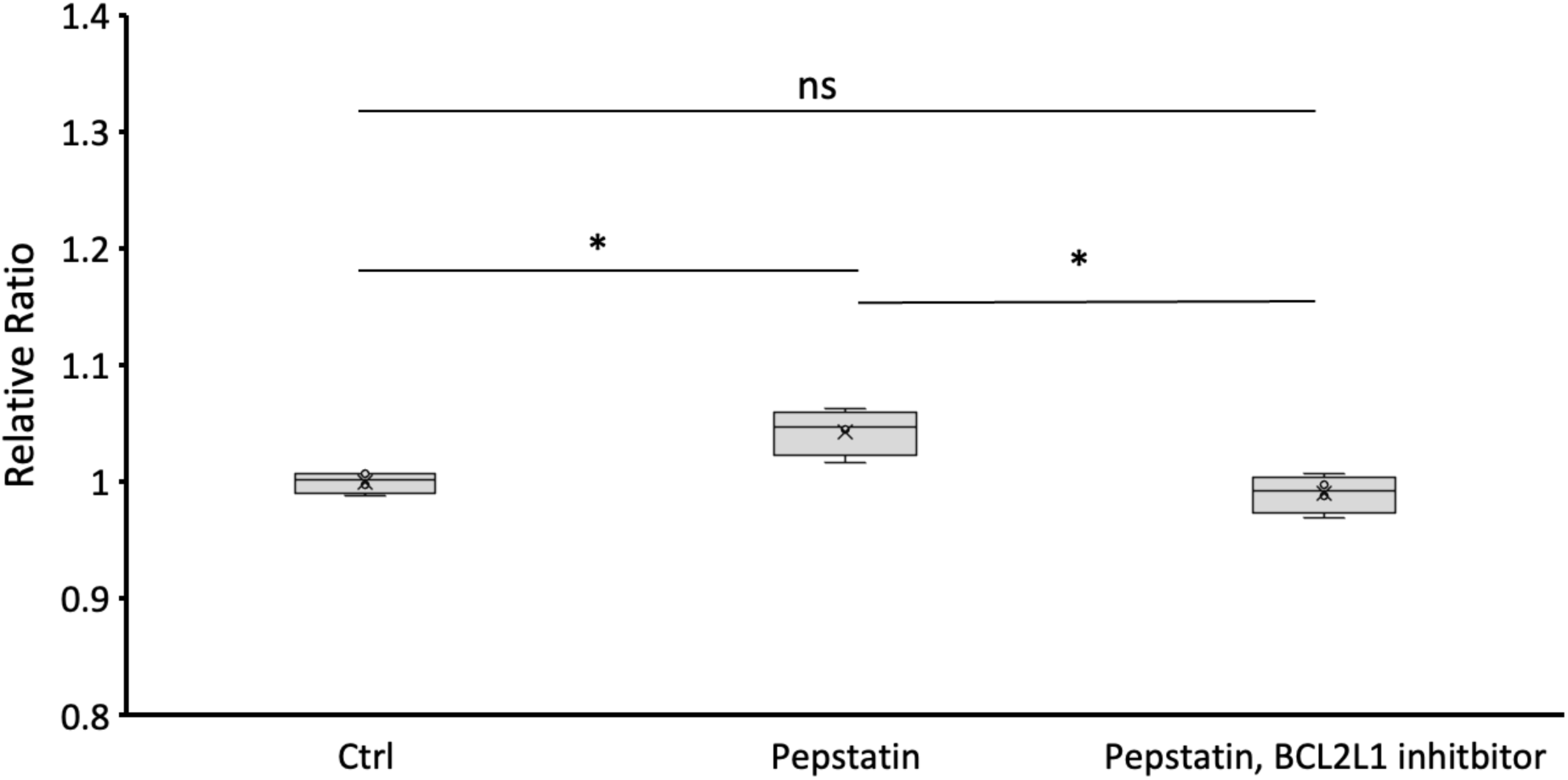
Relative survival rate ratios of TP cells under Control (Ctrl), Pepstatin-treated, and Pepstatin plus BCL2L1 inhibitor co-treated conditions (N = 4). P-values were calculated using Student’s *t*-test. ns (not significant); p < 0.05 *.

**Figure S19.**
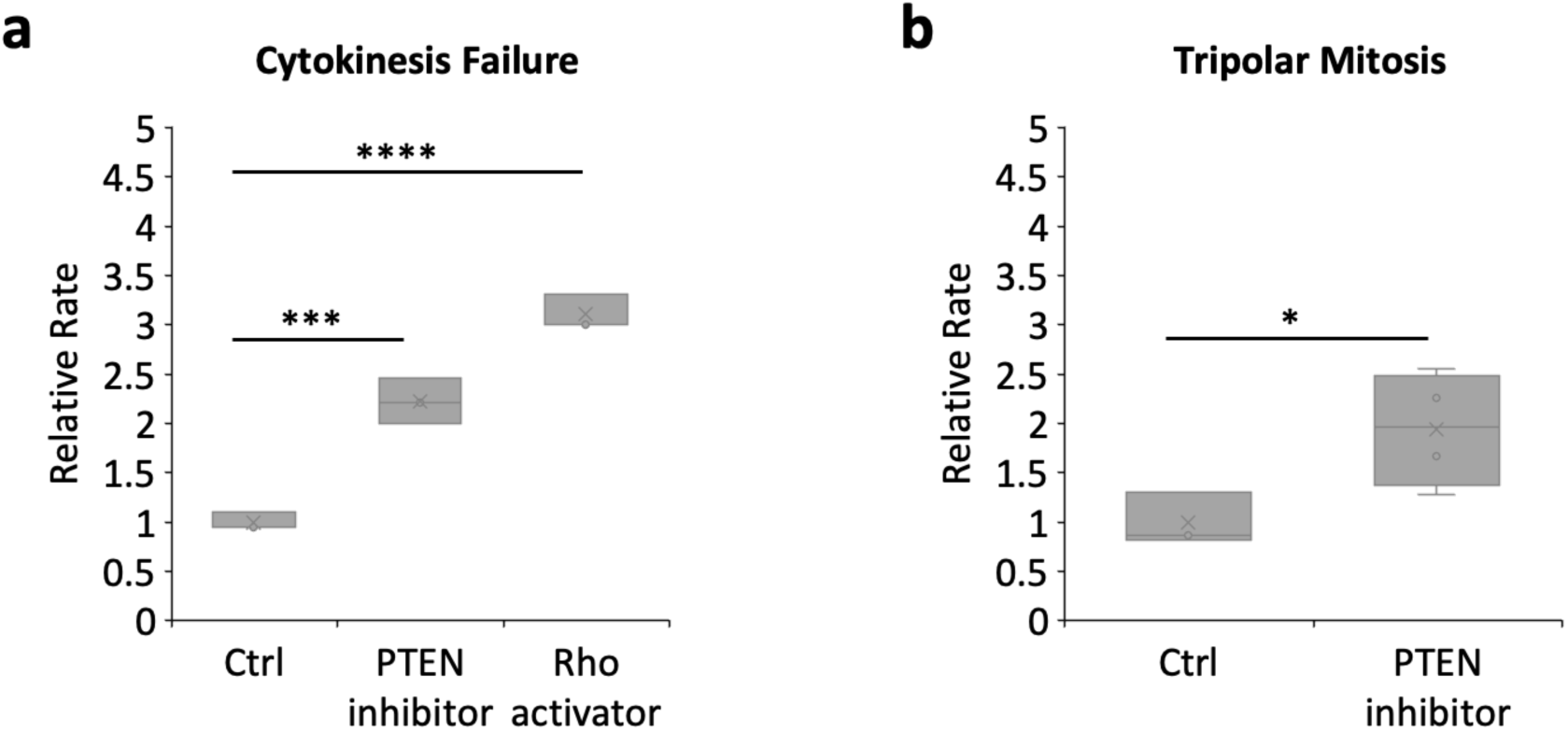
Upregulation of RhoGTPase signaling and downregulation of PTEN in MCF7 cells. a) Statistics of cytokinesis failure rate in MCF7 cells treated without drugs (Control, Ctrl), with a PTEN inhibitor, or with a RhoGTPase activator (N = 3, Student’s t-test). b) Statistics of tripolar mitosis rate in MCF7 cells treated without drugs (Control, Ctrl) or with a PTEN inhibitor (N = 3). The p-value was obtained using Student’s t-test. In the box and whisker plots, the box shows the interquartile range (25th, 50th, 75th percentiles), and whiskers indicate the minimum and maximum values. The p-value was obtained using Student’s t-test. ns (not significant); p < 0.05 *; p < 0.01 **; p < 0.001 ***; p < 0.0001 ****.

